# Stage-specific proteomic signatures of synaptopathy in Alzheimer’s disease

**DOI:** 10.64898/2026.10.02.755976

**Authors:** Tatiana Georgiades, Jessica Griffiths, Fontaine Gibbs, Eileen Chandra, Marianna Papageorgopoulou, Daniel Liang, Harry J. Whitwell, Paul M. Matthews, Samuel J. Barnes, Seth G.N. Grant, Johanna S Jackson

## Abstract

Synapse loss is the strongest pathological correlate of cognitive decline in Alzheimer’s disease, yet the molecular basis of synaptic vulnerability across disease progression remains poorly understood.

We performed proteomic profiling of isolated synaptoneurosomes from *post-mortem* middle temporal gyrus tissue from 67 individuals, comprising 40 individuals spanning all Alzheimer’s disease Braak stages and 27 non-diseased controls. Differential abundance and weighted gene co-expression network analyses were combined with pathway, cell-type and gene set variation analyses to characterise stage-associated changes in the synaptic proteome. Brain-derived candidate proteins were subsequently evaluated using receiver operating characteristic and regression analyses against cognitive examination scores in two independent plasma proteomics datasets.

We identified six distinct stage-associated synaptic proteomic signatures. Mitochondrial function and synaptic signalling declined by mid-stage Alzheimer’s disease, while clathrin-mediated endocytosis increased at mid-stage without further change at later stages. By contrast, proteostasis and perisynaptic astrocytic processes increased progressively throughout the disease course. Global excitatory and inhibitory protein signals also showed a progressive shift in their relative balance (*P* = 7.80 × 10⁻^3^; adjusted *R²* = 0.072). These findings guided targeted investigation of candidate plasma markers linked to distinct synaptic processes in two independent proteomic datasets. We nominated 42 plasma proteins mapping to synaptic signatures of progression, which showed associations with disease stage and cognitive measures including Mini-Mental State Examination, Montreal Cognitive Assessment and Clinical Dementia Rating. PHGDH was prioritised across analyses through associations with cognitive impairment from early disease stages (Clinical Dementia Rating transition 0 to 0.5, false discovery rate = 8.87 × 10⁻^18^), neuropathological progression (*P* = 6.85 × 10⁻^7^) and excitatory-inhibitory protein imbalance (*P* = 5.56 × 10⁻^3^).

Together, our findings define the stage-associated molecular changes at the AD synapse and connect distinct brain pathways to accessible plasma candidates, establishing a mechanistically anchored framework for future patient stratification and evaluation of synapse-directed therapies.

## Introduction

Alzheimer’s disease (AD) is characterised by amyloid beta (Aβ) and tau accumulation, which are central to its neuropathology and diagnosis^1^. However, Aβ burden correlates poorly with cognitive decline, whereas tau shows a stronger yet still modest association^2–4^. Synapse loss, by contrast, is the strongest pathological correlate of cognitive decline^5,6^. Crucially, synaptic dysfunction precedes overt synapse loss, creating a therapeutic window in which compromised synapses may remain amenable to rescue^7^. Existing synaptic markers, such as SV2A PET imaging and CSF proteins neurogranin, SNAP-25 and NPTX2, provide valuable information about synaptic density but remain incompletely characterised with respect to the specific molecular processes they represent^8–11^. Mechanism-linked biomarkers could support earlier identification of synaptic alterations and provide pharmacodynamic readouts of the processes targeted by therapies aimed at restoring synaptic function. Synaptic dysfunction encompasses multiple processes, including shifts in excitatory-inhibitory signalling balance, impaired synaptic plasticity, and dysregulated expression of proteins essential for neurotransmission and memory formation^12–15^. The absence of accessible markers linked to these processes limits the development and clinical evaluation of targeted synaptic therapies.

A thorough understanding of synaptopathy across AD stages requires systematic characterisation of the synaptic proteome. Although alterations in the synaptic proteome have been explored in relation to regional pathology, cognitive status and genetic risk factors^7,15–17^, a characterisation of synapse-specific proteomic changes across neuropathological AD progression is lacking^18^. Furthermore, no framework currently links synaptic proteome alterations to cognitive decline or candidate fluid biomarkers. This represents an important knowledge gap, as mechanistically informed biomarkers associated with earlier disease stages could support, following longitudinal validation, earlier intervention and synapse-directed therapeutic development^19^. Here, we address this gap by mapping the synaptic proteome across AD Braak stages and leveraging the resulting pathway signatures to prioritise novel candidate plasma markers associated with cognitive severity. We also capture changes in excitatory-inhibitory protein balance and perisynaptic astrocytic processes, with PHGDH providing an illustrative candidate connecting several brain and plasma findings. By integrating stage-spanning synaptoneurosome and plasma proteomics, we define stage-associated molecular signatures at the AD synapse and provide a brain-informed framework for nominating candidate plasma markers linked to distinct synaptic processes.

## Materials and methods

### Case selection

We obtained fresh frozen brain tissue samples from seven brain banks in the UK: South West Dementia Brain Bank, London Neurodegenerative Diseases Brain Bank, Multiple Sclerosis and Parkinson’s Tissue Bank, Edinburgh Brain Bank, Oxford Brain Bank, Manchester Brain Bank and Cambridge Brain Bank. Tissue was sampled from the middle temporal gyrus (MTG; BA20/21) of 27 non-diseased control cases (NDC; Braak 0-2), 26 mid-staged AD cases (Braak 3-4) and 14 late-staged AD cases (Braak 5-6). Cases with clinical comorbidities, including diabetes, symptomatic atherosclerotic disease or stroke, or co-pathologies such as Lewy bodies, TDP-43 and vascular disease, were excluded from the cohort. Presence of cerebral amyloid angiopathy or hypertension was not cause for exclusion. Samples that were not previously genotyped for APOE were genotyped in-house. Briefly, DNA was extracted from 15mg of brain tissue using the DNeasy Blood & Tissue Kit (Qiagen) and genotyped via the TaqManTM SNP Genotyping Assay (ThermoFisher) for SNPs rs429358 and rs7412, along with the TaqMan™ Genotyping Master Mix (ThermoFisher), according to the manufacturer’s instructions. All case metadata are recorded in **Supplementary Table 1**.

### Synaptic protein enrichment

Enrichment of synaptoneurosomes was conducted using the Synaptic Protein Extraction Reagent (Syn-PER, Thermofisher), according to the manufacturer’s instructions. 100mg of fresh frozen tissue was suspended in Syn-PER and Complete Protease Inhibitor (EDTA Free, Roche) and homogenised using the SciQuip Basic20 LED Digital Overhead Stirrer. The final pellets were stored at -80°C.

### Immunohistochemistry

Immunohistochemistry staining was performed on paired 8μm Formalin-Fixed, Paraffin-Embedded (FFPE) slides of the MTG region of the cases selected in the study (**Supplementary Table 1**). After slides were deparaffinised in xylene and rehydrated using graded ethanol solutions and endogenous peroxidase activity was blocked with 3% H2O2, followed by antigen retrieval (steamer, EDTA buffer, pH 9.0). Tissue was blocked in 2.5% Normal Horse Serum (MP-7402, Vector Laboratories) in 0.3% PBS-T buffer and then incubated overnight at 4°C with either of the following primary antibodies: 4G8 (800702, 1:10000, Biolegend) and PHF1 (1:5000, The Feinstein Institutes for Medical Research). The ImmPRESS kit (MP-7402, Vector Laboratories) and DAB (SK-4105, Vector Laboratories) were used for antibody visualisation, and Mayer’s haematoxylin solution (Sigma-Aldrich) was used for counterstaining. Stained immunohistochemistry slides were scanned with Leica’s Aperio AT2 DX scanner at 20X magnification. Positive pixel area analysis was performed via HALO software (v3.6, Indica Labs) using antibody-specific optimised macros to quantify immunoreactivity.

### Label-free quantitative proteomics

Proteomics was performed on synaptic samples as described previously^1^. Briefly, dried synaptic pellets were lysed in 8M urea, followed by sonication in a water bath and centrifugation at 4°C, 10,000xg for 10 minutes. Protein concentration was determined using the DC Protein Assay microplate protocol (Bio-Rad), as per the manufacturer’s instructions. Synaptic proteins were subsequently digested and acidified. Digested peptides were desalted using an Oasis HLB 96-well µElution Plate (Waters) and dried in a SpeedVac (Eppendorf) at 60°C for approximately 90 minutes. Peptides were analysed by liquid chromatography-tandem mass spectrometry (LC-MS/MS) in three separate batches: the first batch included NDC and Braak 5-6 AD cases, while the two other batches comprised NDC, Braak 3-4 and 5-6 AD cases. Processing of samples was conducted using the ACQUITY UPLC M-Class system (Waters) coupled with a ZenoTOF 7600 system (Sciex) in Zeno SWATH Data Independent Acquisition mode.

### Processing of proteomics data

Raw data was processed using DIA-NN (v1.9) and mapped to a predicted library of canonical human proteins from Uniprot. All batches were searched with a Uniprot version downloaded on 07/02/2025. All further processing was conducted in R version 4.4.1. Proteins were filtered for 30% missingness and raw abundance values were log2-transformed. Data was normalised using median centring and one sample was excluded due to 100% missingness upon filtering. Robust sequential imputation was conducted on the data using impSeqRob from the MSnbase v2.32.0 package^2^. We accounted for batch effects by adding batch as a covariate in all models.

### Differential expression analysis

Differential protein expression between NDC and AD groups as well as moderate and late-stage AD was conducted using limma package version 3.62.1^3^. Due to low statistical power to detect late compared to mid-stage AD changes, the mid-stage disease signal was lowered in the contrast design by averaging NDC with AD Braak 3-4. We applied empirical Bayes moderation and *P*-values were corrected for multiple testing via the Benjamini-Hochberg (BH) method to control the false discovery rate (FDR)^4^. All differential expression models included age, sex, post-mortem interval (PMI) and batch as covariates. We evaluated the impact of covariates on effect sizes of differentially expressed proteins (DEPs) by applying the same models without adjusting for covariates and performing Pearson correlation on the effect sizes of the adjusted compared to unadjusted models (**Supplementary Fig. 1**). Additionally, we evaluated the Pearson correlation of DEPs with individual batches and Braak stages. All DEPs showing more significant Pearson correlation with technical batch than with pathological disease stage were excluded. All ambiguously mapped synaptic DEPs were also excluded, and the remaining proteins were reported.

### Weighted Gene Co-expression Network Analysis

Weighted Gene Co-expression Network Analysis was conducted using the WGCNA (v1.73) on the upper 75^th^ percentile of proteins with highest variance across samples to remove noise^5^. Hierarchical clustering of samples was used to detect potential outliers. A signed hybrid protein co-expression network was constructed using biweight midcorrelation to calculate positive correlations of proteins in a robust method against outliers. A soft thresholding power of 9 was selected as the lowest threshold to produce scale free topology model fit correlation 0.85, upon testing powers from 1 to 20 (**Supplementary Fig. 3C**). A protein clustering tree was constructed via hierarchical clustering using topology matrix dissimilarity and cut using the cutreeDynamic() function with minimum cluster size of 30 proteins and parameter deepSplit=4 to ensure small interpretable cluster sizes. Upon cutting the protein clustering tree at a default height of 0.998, 14 modules were produced and an additional ‘unassigned’ module (M0/grey), including proteins that did not correlate with any of the modules. Module membership of each protein was derived by computing the kME value. Module-trait correlations were calculated via Pearson correlations of module eigenproteins with traits, including Braak stage, age at death, sex, post-mortem interval (PMI), technical batch, amyloid (4G8) and pTau (PHF1) percentage area stained covered in the MTG. Correlation of module eigenproteins with Braak stage was also validated upon correction for covariates age, sex, PMI and batch, via linear regression. All *P*-values were FDR adjusted via the BH correction method for multiple testing.

### Pathway and cell-type enrichment analysis

To prioritise the most representative proteins of each module for pathway enrichment analysis, proteins per module were filtered to remove the lower 10^th^ percentile of proteins by kME. Pathway enrichment analysis was conducted per module using the enrichR (v3.4) package^6^. The following pathway databases were used: GO Biological Process 2025, GO Cellular Component 2025, KEGG 2021 Human, Reactome Pathways 2024 and SynGO 2024. The total synaptic proteins detected upon processing of label-free quantitative proteomics was used as background in the enrichment analysis. Gene ratio was derived using the count of enriched genes against the background genes in the pathway. Upon filtering for significantly enriched pathways by FDR, we calculated the association of each pathway with Braak stage, to facilitate with prioritisation of the most relevant pathways to disease progression. In short, individual protein correlations to Braak stage were computed using a linear regression model adjusted for age, sex, PMI and batch. We used Fisher’s method to combine *P*-values of genes enriched in each pathway to produce a single *P*-value per pathway for association with Braak. Fisher’s *P*-values were adjusted via BH for multiple testing. Pathways were filtered for significant association with Braak and prioritised based on pathway effect size, calculated as the mean Braak-stage regression coefficient across represented proteins. For the summary heatmap, representative significant pathways were selected across databases to capture the principal functional themes of each module and grouped into broader biological categories.

Expression weighted cell-type enrichment was performed via the EWCE (v1.14.0) package^7^. We used a reference dataset of cell types in the human MTG, from the Allen Human Brain Atlas^8^. We used all proteins included in each module to identify all possible enriched cell types. *P*-values were adjusted via the BH method.

### Gene Set Variation Analysis

We performed Gene Set Variation Analysis using the GSVA (v2.0.0) package^9^. We used module proteins as gene sets to calculate the per sample enrichment of each module that is significantly associated with Braak stage. Due to the nature of protein abundance data, Gaussian kernel was used for non-parametric estimation of the empirical cumulative distribution function of expression levels across samples. GSVA scores per module, were subsequently used for differential pathway analysis using limma to determine module expression changes across AD stages. Similarly to differential protein expression analysis, models were adjusted for age, sex, PMI and batch. Contrasts were set to compare Braak 3-4 to Braak 0-2, Braak 5-6 to Braak 0-2 and Late to Moderate AD (Braak 5-6 to averaged Braak 3-4 and 0-2). Empirical Bayes moderation was applied, and *P*-values were corrected for multiple testing via BH.

### Excitatory – Inhibitory protein balance

To derive a signature of excitatory-inhibitory (E-I) protein balance in the synaptic proteomics, we initially identified all excitatory and inhibitory proteins present in the dataset, as defined by Lauterborn et al., 2021^10^. We firstly calculated a per sample excitatory signal using mean z-scored abundances of excitatory proteins in our dataset and repeated this process with inhibitory proteins to calculate an inhibitory signal. For the E-I protein balance signature, the inhibitory signal was subtracted from the excitatory signal for each sample. Linear regression adjusted for age, sex, PMI and batch was conducted to capture the monotonic change of E-I protein balance across Braak stages.

### Independent dataset analysis

We used the Global Neurodegeneration Proteomics Consortium (GNPC) dataset for orthogonal validation^11^. We selected all cases with an AD diagnosis and plasma proteomics, excluding cases with multiple diagnoses, missing sex and age at visit. For selection of controls, we included all recruited control cases or cases without a diagnosis of any neurodegenerative disease and clinical dementia rating (CDR) 0. We also excluded any cases with mild cognitive impairment (MCI). We used SomaScan 7k assay proteomic data from citrate and EDTA plasma from each case at the latest age at visit. In total, we used 2309 cases with AD and 2418 controls (**Supplementary Table 2**). We conducted all proteomic data processing and differential expression analysis as described in Ali et al., 2025^12^. GNPC data were analysed in AD Workbench, as part of the AD Data Initiative, in R v4.5.2. We used z-scored aptamer expression in all analyses with this dataset, as it was required for the harmonisation of multiple cohorts. The differential expression model was adjusted for age at visit, sex, PC1 and PC2. We also performed independent validation using the SomaScan plasma proteomics data and related list of DEPs from Dammer et al., 2022^13^. All analyses with this dataset were performed on batch corrected log-transformed aptamer expression values, in R v4.4.1.

Synaptic DEPs were filtered to be present in the GNPC, Dammer et al., 2022 and Ali et al., 2025 datasets^12,13^. We produced Receiver Operating Characteristic (ROC) curves using the pROC (v1.19.0.1) package. We used Area Under the Curve (AUC) and bootstrapped (*n*=1000) 95% confidence intervals (CI) to evaluate the overall performance of each protein (**Supplementary Tables 3-6**). The GNPC dataset was used to examine discrimination between AD/Control and different stages of AD. To adjust for the effect of different cohorts, models were adjusted for cohort site. No other covariates were added, to evaluate the discriminative performance of proteins. The Dammer et al., 2022 dataset was used solely for AD/Control distinction, due to power constraints. AD stages were determined based on cognitive examination scores as shown in **Supplementary Table 7**, based on cut-offs set in the literature^14,15^.

In disease-stage ROC analyses comparison groups were downsampled to the size of the group with the lowest sample size, to avoid sample size-based bias. Sensitivity and specificity were calculated for the optimal cut-off, as well as Youden’s index as a metric to evaluate the discriminative performance of proteins.

Linear regression of candidate biomarker protein aptamers was conducted with Mini-Mental State Examination (MMSE) and Montreal Cognitive Assessment (MoCA) scores as outcome variables. In the GNPC dataset, models were adjusted for age at visit, sex, PC1 and PC2. The Dammer et al., 2022 dataset was used solely for regression analysis with MoCA scores, where the linear model was adjusted for age at visit and sex. All *P*-values were corrected for multiple testing via the BH method. Moreover, Clinical Dementia Rating (CDR) scores from the GNPC dataset were modelled against aptamer expression corrected for age at visit, sex, PC1 and PC2, via ordinal logistic regression, using the ordinal (v2023.12.4.1) package^16^. Nominal test was performed to identify potential violations of the proportional odds assumption in each protein model. Where the proportional odds assumption was violated due to non-linear change across CDR scores, a partial proportional odds model was applied. For each model the odds ratio of being in an increased CDR score and the standard error was calculated. When a partial proportional odds model was applied, statistics were calculated for the optimal threshold of CDR score change.

### Immunofluorescence

Formalin-fixed paraffin-embedded tissue sections were heated at 60°C for 3h, dewaxed twice in xylene and once in 100% ethanol for 5min each. Sections were rehydrated sequentially in 100%, 90% and 70% ethanol followed by distilled water, with each step performed for 5min. For antigen retrieval, slides were immersed in preheated 1 x EDTA buffer (pH 9) and steamed for 20min. After cooling to room temperature slides were washed for 5 min each in distilled water and 1x phosphate-buffered saline (PBS; 11510546, Gibco™).

Tissue sections were outlined using a hydrophobic barrier pen (ab2601, Abcam) under a humidified chamber. Sections were blocked for 1h at room temperature with 200μL of 10% normal donkey serum (NDS; ab7475, Abcam) prepared in 0.3% PBS-Triton X-100. Primary antibodies (**Table 1**) were diluted to the indicated concentrations in the same blocking solution, applied at 200μL per section and incubated overnight at 4°C. Primary antibody incubation was followed by 150μL of 1x TrueBlack Lipofuscin Autofluorescence Quencher (23007, Biotium), for no longer than 1min. Slides were then washed three times for 5min in 1x PBS. Secondary antibodies (**Table 1**) were diluted 1:1000 in 1x PBS, applied at 200μL per section and incubated for 1h at room temperature in the dark. Sections were subsequently washed three times for 5min in 1x PBS.

**Table 1:** Primary and secondary antibody information for immunofluorescence staining of FFPE sections. Details include host species, dilution factor, catalogue number and secondary antibodies used for primary antibody detection. AF, Alexa Fluor; FFPE, Formalin-Fixed Paraffin Embedded.

| Primary Antibody | Species | Dilution | Catalogue No. | Secondary Antibody |
| --- | --- | --- | --- | --- |
| PHGDH | Mouse | 1:200 | MA5-31357<br>(ThermoFisher) | DxM AF647<br>(A-31571,<br>Invitrogen) |
| Ezrin | Rabbit | 1:500 | 16814585<br>(ThermoFisher) | DxRb AF568<br>(A10042,<br>Invitrogen) |
| Synaptophysin | Guinea Pig | 1:500 | 101 308<br>(Synaptic<br>Systems) | DxGP AF488<br>(706-545-148,<br>Jackson) |

Nuclei were counterstained for 10min with DAPI (1:1000 in 1x PBS, 62248, Thermo Scientific™; 200μL per section), followed by a final 5-min wash in 1x PBS. Coverslips were mounted using 15μL of ProLong Diamond Antifade Mountant (P36970, Invitrogen) and slides cured for 24h before imaging.

### Confocal Imaging

Slides were imaged using the Leica Stellaris microscope (Leica Microsystems). Images were acquired using a 63x objective with immersion medium (11944399, Leica Microsystems). Z-stacks were collected at 0.5-μm intervals. Acquisition parameters, including the laser lines and laser powers, were held constant across all samples. Images were acquired from three regions of interest per case in the grey matter. The experimenter remained blinded to disease status during image acquisition.

### Image analysis and statistical comparison

Image processing and quantitative analysis were performed using Imaris v11.0.0 (Oxford Instruments). The experimenter remained blinded to disease status during image analysis. Gaussian background subtraction was applied, followed by adjustment of image brightness and contrast to facilitate visualisation.

Synaptophysin^+^ puncta and PHGDH–ezrin colocalised signal were segmented using the Imaris Surfaces tool. Synaptophysin surfaces were generated using local-contrast background subtraction, with touching objects separated by morphological splitting. Segmented surfaces were subsequently filtered and manually reviewed to exclude autofluorescent structures that were not consistent with the target signal.

Spatial association between synaptophysin and PHGDH–ezrin was assessed using the shortest distance between the synaptophysin components and the PHGDH–ezrin surface. Synaptophysin components with shortest distance of ≤0μm were classified as colocalised. The percentage colocalisation for each image was calculated by dividing the number of colocalised synaptophysin components by the total number of synaptophysin disconnected components and multiplying by 100.

Synaptophysin puncta density was calculated by dividing the total number of disconnected synaptophysin components by the physical volume of the corresponding image stack and was expressed as puncta per μm³. PHGDH–ezrin signal was quantified as the summed volume of all segmented PHGDH–ezrin surface components divided by the total stack volume and expressed as a percentage. Measurements were calculated separately for each stack and averaged across the included stacks from each case, such that each stack contributed equally to the case-level estimate. Kruskal-Wallis followed by Dunn’s post-hoc tests and BH correction was used for group comparisons. Monotonic associations with ordered neuropathological stage were assessed at the case level using Spearman’s rank correlation, with *P* values BH-adjusted across imaging outcomes. All data analysis was conducted using R 4.4.1.

## Results

### Synaptic protein alterations increase with advancing AD neuropathology

We used *post-mortem* human brain tissue from the middle temporal gyrus (MTG) region of 26 AD cases with mid-stage (Braak 3-4) and 14 cases with advanced pathology (Braak 5-6), alongside 27 non-diseased controls (NDC, Braak 0-2) and processed this tissue to enrich for synaptoneurosomes (**Supplementary Table 1**). Synaptic proteomics was conducted on the synaptoneurosome samples, and a total of 5300 proteins were identified with 4035 across all samples after quality control.

We performed differential expression analysis (|Log2FC| > 0.5, FDR < 0.05) to compare samples from mid-stage and advanced AD with NDC samples (**Fig. 1A-C, Supplementary Table 8**). We identified 148 differentially expressed proteins (DEPs; 82 upregulated, 66 downregulated) between mid-stage AD and NDC (**Fig. 1A**), and 282 DEPs between advanced AD and NDC (196 upregulated and 86 downregulated; **Fig. 1B**). To derive a late-stage proteomic signature, we compared late with moderate AD cases (average of Braak 3-4 and NDC groups). We detected 60 DEPs (46 upregulated and 14 downregulated; **Fig. 1C**), which reflect changes occurring at late pathological stages of AD. These results indicate differential synaptic protein changes across the pathological progression of AD, with advanced AD pathology associated with a greater number of synaptic proteomic alterations than mid-stage AD.

**Fig. 1:**
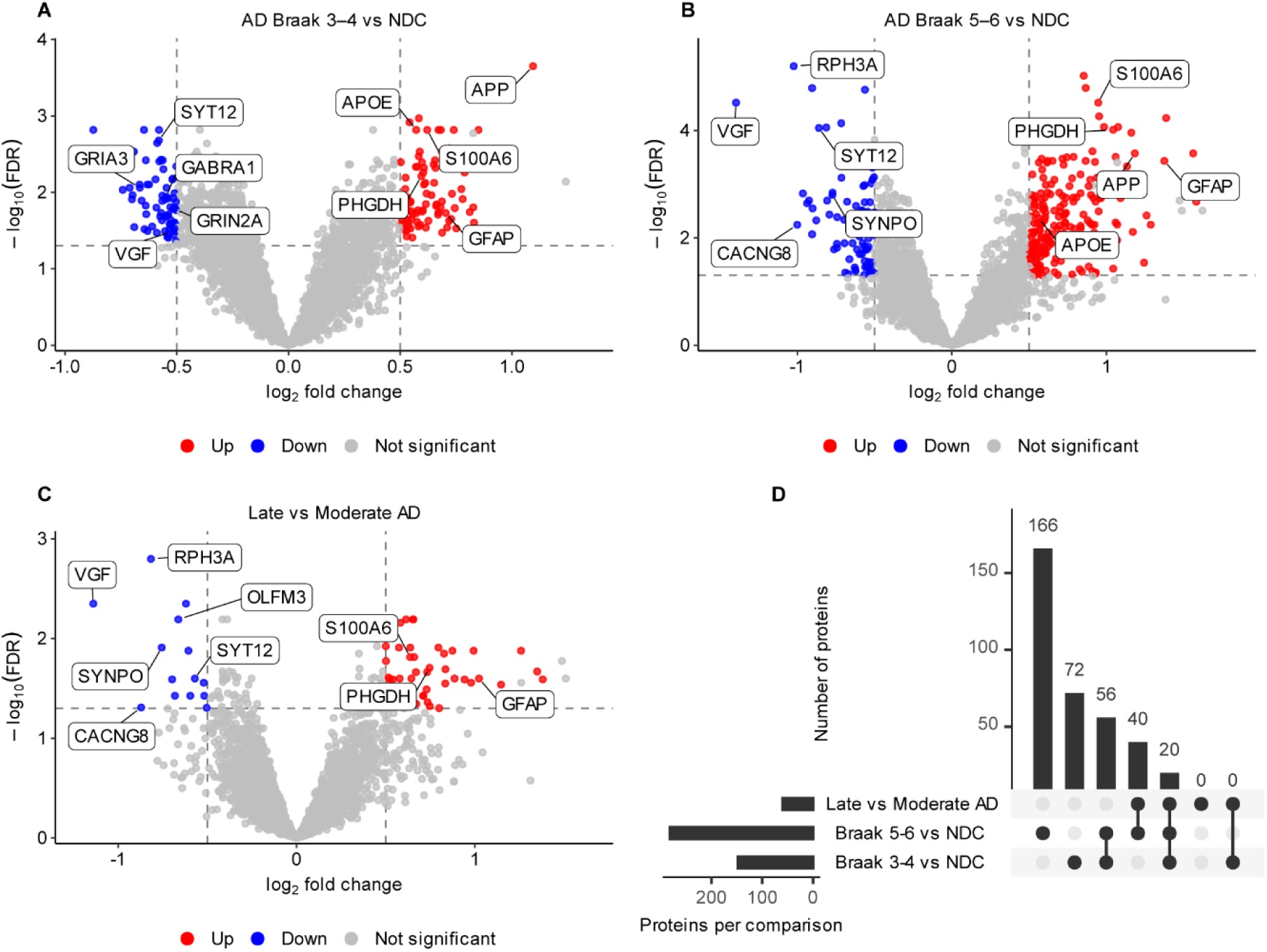
Synaptic protein alterations increase with advancing AD neuropathology. (**A-C**) Volcano plots of differential expression analysis for (**A**) mid-stage AD (Braak 3-4, *n* = 26) vs NDC (Braak 0-2, *n* = 27) (**B**) late-stage AD (Braak 5-6, *n* = 14) vs NDC (**C**) Late (Braak 5-6) compared to moderate AD (average of Braak 3-4 and 0-2). Linear model adjusted for age, sex, *post-mortem* interval and batch. FDR < 0.05, |Log2FC| > 0.5. Known DEPs involved in AD or synapses are labelled. Grey points passing the FDR and Log2FC thresholds denote ambiguously mapped proteins. (**D**) Upset plot showing overlap among filtered DEPs across the three comparisons. *P* values were adjusted using the Benjamini–Hochberg method. AD, Alzheimer’s disease; NDC, non-diseased control; DEP, differentially expressed protein; FDR, false discovery rate.

We examined the overlap of DEPs across AD stage comparisons to determine their specificity to different stages of disease progression (**Fig. 1D**). Seventy-two DEPs were detected only in mid-stage AD vs NDC. Of the 206 proteins detected in late but not mid-stage AD, 166 were identified only in late-stage AD vs NDC, whereas 40 were also identified in the Late vs Moderate AD comparison. Seventy-six DEPs were shared between mid-stage and late-stage AD relative to NDC. Of these, 56 were not detected in the Late vs Moderate AD comparison, whereas 20 were significant across all three comparisons and showed concordant directionality. Thus, all 60 Late vs Moderate AD DEPs were also detected in late-stage AD vs NDC: 40 were specific to late-stage AD, whereas 20 were already altered at mid-stage and continued to change in the same direction, suggesting that relatively few individual proteins change consistently across all stages.

Based on these patterns, we classified the DEPs as Mid-stage AD (72 proteins), Late AD (206 proteins) or AD/NDC (76 proteins), with the 20 proteins significant across all three comparisons additionally classified as Progression proteins (**Supplementary Table 9, Supplementary Fig. 2**). Mid-stage AD changes included reductions in proteins involved in GABAergic and glutamatergic signalling (GABRA1, GRIN2A), whereas Late AD changes included reductions in proteins involved in synaptic vesicle release and postsynaptic organisation (RPH3A, CACNG8, SYNPO). The AD/NDC signature included increases in proteins canonically linked to AD (APP, APOE). Progression proteins included reductions in proteins linked to synaptic signalling and plasticity (SYT12, VGF) and increases in astrocyte-associated proteins (GFAP, S100A6, PHGDH). This classification identified candidate proteins associated with distinct stages or trajectories of AD pathology and provided the foundation for subsequent plasma analyses.

### Distinct synaptic signatures are associated with pathological AD progression

To resolve coordinated synaptic protein changes occurring across AD progression, we identified signatures of co-expressed proteins associated with Braak stage using Weighted Gene Co-Expression Network Analysis (WGCNA; **Fig. 2**). A co-expression network was constructed on proteins with 75% highest variance across samples according to a previous protocol^17^ (**Supplementary Fig. 3A**). Based on these parameters we derived distinct modules of co-expressed synaptic proteins (**Supplementary Fig. 3D-E**). We identified modules associated with progression of AD pathology by calculating the Pearson correlation of module eigenproteins with sample traits (**Fig. 2A**). We detected six modules (M1, M2, M4, M6, M7 and M13) significantly associated with Braak stage. Of the six progression-associated modules, four modules displayed increasing (M4, M6, M7, M13) and two showed decreasing (M1, M2) eigenprotein expression across Braak stages (**Fig. 2B**). Overall, these modules reflect synaptic protein co-expression changes associated with AD progression, that may be differentially driven by amyloid or pTau.

**Fig. 2:**
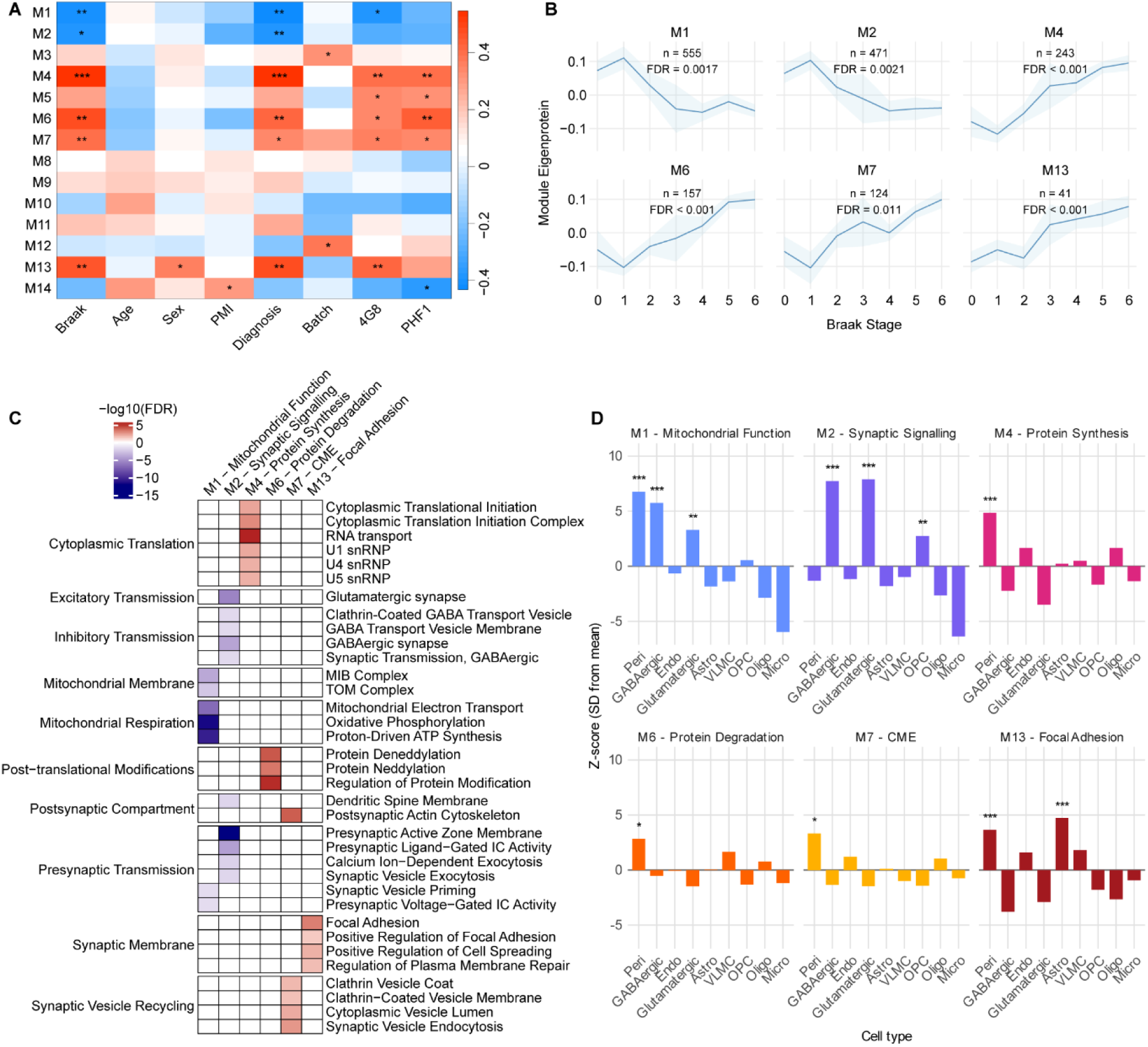
Distinct synaptic proteomic signatures are associated with AD progression. (**A**) Module eigenprotein Pearson correlation matrix with sample traits. (**B**) Module eigenproteins significantly correlated with Braak stage in linear regression adjusted for age, sex, *post-mortem* interval and batch. The line and shaded band represent mean module eigenprotein across Braak and SEM. FDR < 0.05. *n*= number of proteins included in the module. (**C**) Pathway enrichment analysis heatmap from four pathway databases on the top 90 percentile proteins by module membership *kME* per module. Significantly enriched pathways were prioritised based on pathway significance with Braak stage (Fisher’s FDR < 0.05) and effect size. Signed – log10(FDR) was derived from pathway enrichment significance directed by effect size. D) Cell type enrichment analysis was conducted using a human middle temporal gyrus reference from the Allen Human Brain Atlas. BH-adjusted *P*-value < 0.05. * FDR<0.05, ** FDR<0.01, *** FDR<0.001. SEM, Standard Error of the Mean; FDR, false discovery rate.

We identified distinct roles that indicate potential functions of the progression-significant modules via pathway enrichment analysis of proteins with the highest module membership (**Fig. 2C**). In larger modules, significantly enriched pathways were prioritised based on correlation with Braak stage, and gene ratio. Modules were assigned a main functional characterisation based on prevalence of pathways across databases. M1 was primarily associated with mitochondrial function, M2 with synaptic signalling, M4 with protein synthesis, M6 with protein degradation, M7 with clathrin-mediated endocytosis (CME) likely powered by actin polymerisation^18^, and M13 with focal adhesion.

We conducted cell-type enrichment analysis on the progression-associated modules, using a human MTG reference dataset^8^, to identify enrichment of potential non-neuronal interactors (**Fig. 2D, Supplementary Table 10**). M1 – mitochondrial function and M2 – synaptic signalling modules were mainly associated with GABAergic and glutamatergic neurons. Interestingly, M13 was enriched for astrocytic proteins, specifically astrocytic markers previously associated with astrogliosis in AD pathology, such as GFAP, PBXIP1 and DTNA^19–21^.

We next investigated the association of synaptic signatures with different stages of AD via gene set variation analysis (GSVA). After calculating the per sample enrichment of each module, we conducted differential expression analysis of modules across grouped Braak stages (**Fig. 3A-C**). We compared mid-stage and late-stage AD cases with NDC and late with moderate AD, to determine whether synaptic signatures significantly change across disease progression. Both mitochondrial function and synaptic signalling, demonstrate significant downregulation in earlier stages of AD which remains at later stages (**Fig. 3A-B**), with no further decrease between moderate and late AD (**Fig. 3C**). In addition, CME is upregulated in mid- and late-stage AD compared with NDC but does not show a significant change after moderate AD stages. In contrast, expression of protein synthesis, protein degradation and focal adhesion, increases across all three comparisons (**Fig. 3A-C**). Protein degradation and protein synthesis show the largest differences between late-stage AD and NDC (*Log2FC*=0.80 and 0.78, respectively), followed by focal adhesion (*Log2FC*=0.75; **Fig. 3B**). Protein synthesis displays increases of similar magnitude in earlier and later AD stages (*Log2FC*=0.49 and 0.54). By contrast, the effect size for protein degradation is larger in the Late vs Moderate AD comparison (*Log2FC*=0.63) than in mid-stage AD vs NDC (*Log2FC*=0.34). Focal adhesion/astrogliosis shows a similar, although less pronounced, difference between these comparisons *(Log2FC*=0.56 and 0.39, respectively; **Fig. 3A, C**).

**Fig. 3:**
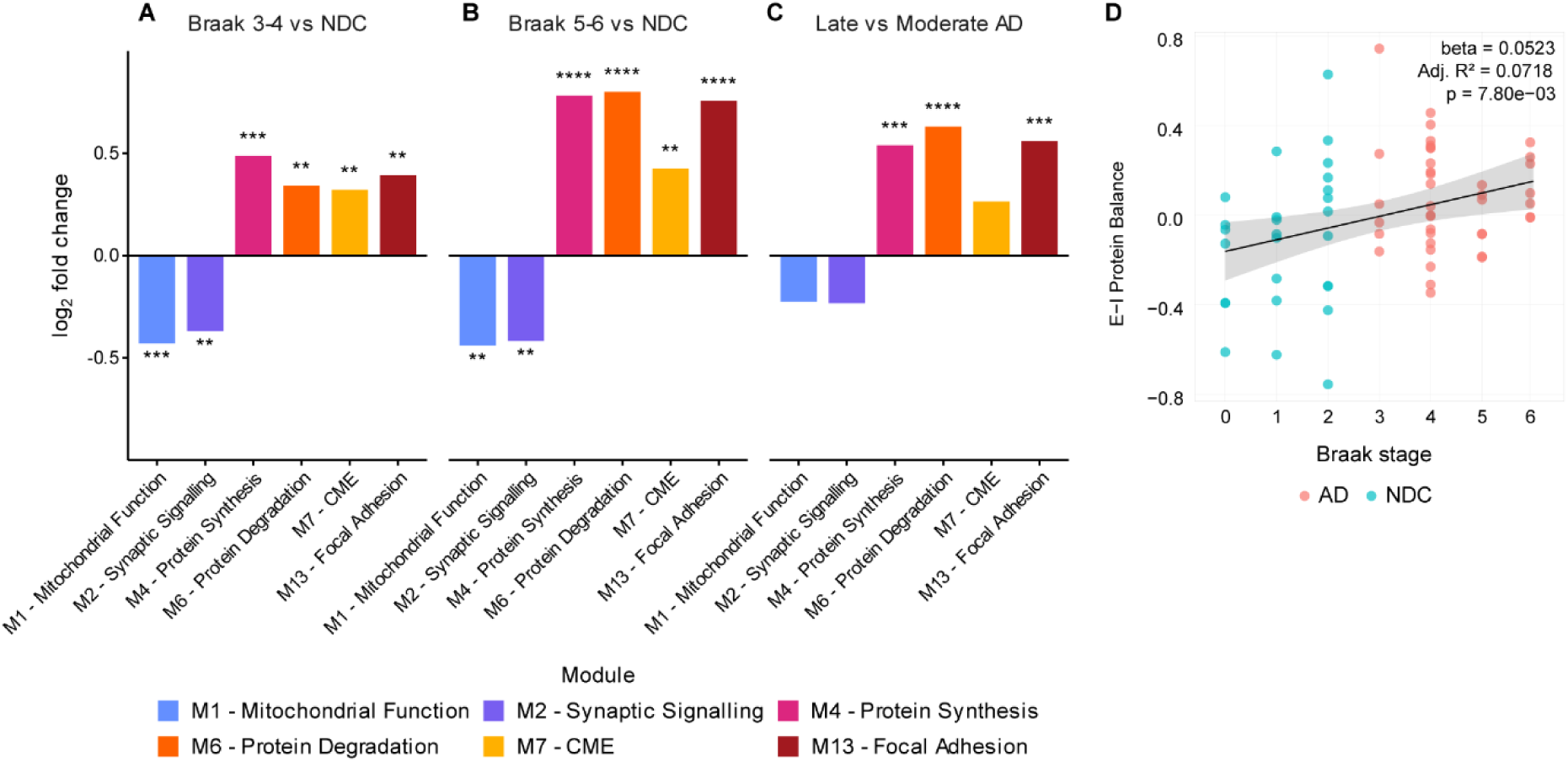
Synaptic signatures follow distinct trajectories across AD stages. (**A-C**) Log2 fold-changes in progression module GSVA scores from differential gene set expression analysis between (**A**) Braak 3-4 vs NDC, (**B**) Braak 5-6 vs NDC and (**C**) Late vs Moderate AD. Linear model adjusted for age, sex, PMI and batch. (**D**) Linear regression of E-I protein balance against Braak stage in synaptic proteomics. E-I protein balance was calculated as the difference between mean z-scored excitatory to inhibitory protein expression. Excitatory and inhibitory proteins were defined using a dataset by Lauterborn et al. Model adjusted for age, sex, PMI and batch. Points are coloured by diagnosis. The line and shaded band represent the fitted linear association and 95% confidence interval. *P* values were adjusted using the BH method. * FDR<0.05, ** FDR<0.01, *** FDR<0.001, **** FDR < 0.0001. NDC, non-diseased control; PMI, post-mortem interval; E-I, excitatory-inhibitory; FDR, false discovery rate.

The enrichment of glutamatergic and GABAergic neuronal proteins in the downregulated modules raised the question of whether excitatory and inhibitory components changed proportionately across AD progression. A proteomic signature of excitatory and inhibitory expression changes was derived using normalised z-scored abundances of previously characterised^10^ excitatory and inhibitory proteins per sample (**Supplementary Fig. 4A-B**). Both the excitatory (**Supplementary Fig. 4A**) and inhibitory (**Supplementary Fig. 4B**) protein signals significantly declined across Braak stages. Interestingly, the magnitude of the linear regression coefficient for the excitatory signal (*β*=-0.084) is lower than the inhibitory signal (*β*=-0.136), pointing to greater reduction in abundance of inhibitory proteins across Braak stage. To examine the relative changes of the two signals directly, we computed a signature of excitatory-inhibitory (E-I) protein balance by subtracting the inhibitory from the excitatory sample scores across Braak stages (**Fig. 3D**). E-I protein balance increased significantly with Braak stage (linear regression, *P*=7.80×10^-3^, Adj *R^2^*=0.072), indicating a progressive shift towards greater excitatory relative to inhibitory protein abundance. This proteomic shift is consistent with previous electrophysiological evidence of synaptic hyperexcitability^22^.

Taken together, these results demonstrate distinct signatures of synaptic processes across AD stages, which provide a biological framework for associating plasma candidates with distinct components of AD-associated synaptic pathology.

### Brain-derived synaptic signatures nominate a mechanistically diverse set of plasma candidates

The stage-associated synaptic proteins identified in our study include proteins characterised in previous studies as well as novel findings. Of 354 unique candidate proteins, only 10 had previously been proposed as synaptic biomarkers, with studies focused predominantly on CSF and late-stage disease (**Supplementary Table 11**). Plasma-accessible markers linked to AD-associated synaptic alterations therefore remain largely unexplored, particularly during earlier disease stages.

We evaluated the candidate markers derived from synaptic proteomics (**Supplementary Table 9**) in two independent SomaScan plasma proteomic datasets: The Global Neurodegeneration Proteomics Consortium (GNPC; **Supplementary Table 2**)^11^ and Dammer et al., 2022^13^. We first identified previously published AD vs Control DEPs in GNPC^12^, and Dammer et al.^13^ (**Fig. 4A**). To validate these findings, we conducted differential expression analysis in the GNPC dataset as previously described^12^, using AD and Control cases with cognitive examination scores concordant with their clinical diagnosis (**Supplementary Fig. 5A**). We then compared DEPs identified in our analysis with those reported in the published datasets (**Fig. 4A, Supplementary Fig. 5B**). After filtering overlapping DEPs against our list of synaptic candidate biomarkers, 51 proteins were identified for further evaluation (**Fig. 4A**; 8/72 (11.1%) Mid-stage AD candidates, 32/206 (15.5%) Late AD candidates, 11/76 (14.5%) AD vs NDC candidates, 4/20 (20.0%) Progression candidates).

**Fig. 4:**
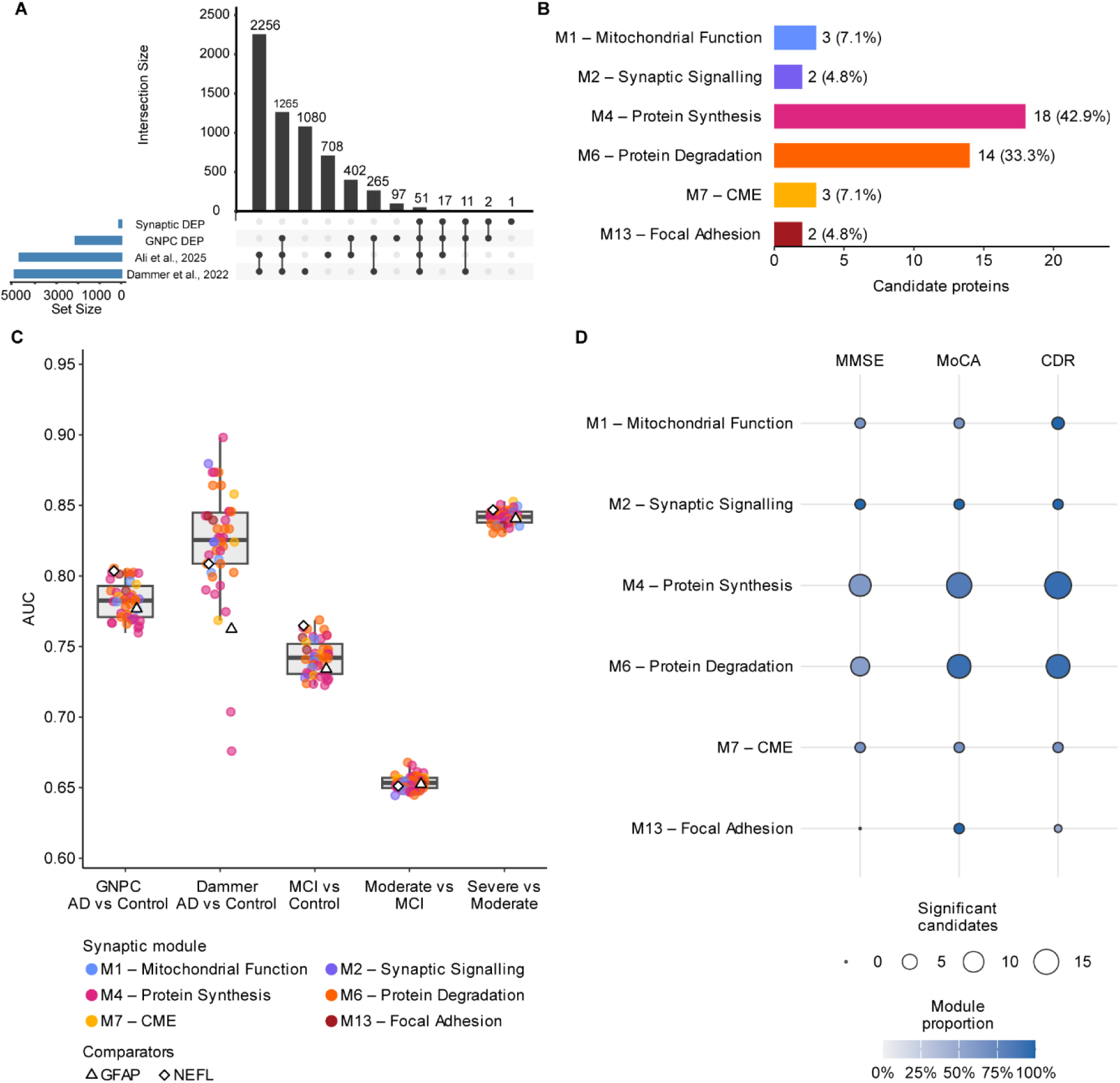
Plasma candidate markers are linked to brain synaptic signatures. (**A**) UpSet plot showing overlap between synaptic DEPs and DEPs from three plasma analyses prioritises 51 candidate proteins. (**B**) Distribution of the 42 candidates mapped to the six Braak-associated synaptic modules. Labels show protein counts and percentages of the 42 mapped candidates. (**C**) AUC distributions for the mapped candidates in GNPC AD versus control, Dammer et al. AD vs Control, and GNPC adjacent cognitive-stage comparisons. Points are coloured by synaptic module; GFAP and NEFL are shown as standard neurodegeneration-associated biomarker comparators. Boxes show the median and interquartile range, with whiskers extending to 1.5 times the interquartile range. (**D**) Significant associations with MMSE, MoCA and CDR by synaptic module. Bubble size represents the number of significant candidates and colour represents their proportion within each module. DEPs, differentially expressed proteins; AD, Alzheimer’s disease; AUC, area under the receiver operating characteristic curve; GNPC, Global Neurodegeneration Proteomics Consortium; MMSE, Mini-Mental State Examination; MoCA, Montreal Cognitive Assessment; CDR, Clinical Dementia Rating.

Of these 51 proteins, 42 (82.4%) mapped to one of the six Braak-associated synaptic modules (**Fig. 4B**). Most candidates mapped to signatures of synaptic proteostasis with 18 assigned to protein synthesis (42.9%) and 14 to protein degradation (33.3%). Three candidates mapped to each of mitochondrial function and CME (7.1% each), and 2 to each of synaptic signalling and focal adhesion (4.8% each). Although concentrated in proteostasis-related modules, the candidates spanned all six signatures, linking plasma-accessible proteins to distinct components of AD-associated synaptic biology.

We first investigated the ability of candidate plasma markers to discriminate AD from Control participants using receiver operating characteristic (ROC) analysis in the two datasets (**Fig. 4C; Supplementary Tables 3-5**). Significant candidates were represented across all six synaptic modules in both datasets, although their discriminatory performance varied (Area Under Curve [AUC]=0.68–0.90). Across the two datasets, 41 of the 42 mapped candidates achieved an AUC within or above the range observed for the established neurodegeneration-associated markers NEFL and GFAP in one or both datasets (AUC=0.76-0.81; **Supplementary Fig. 6A-D**). Although the top-ranked candidate differed between datasets (MVD in GNPC and CSRP1 in Dammer), CNDP2, a Late AD candidate mapping to the protein degradation module, ranked among the leading candidates in both (AUCs=0.80 and 0.86; **Supplementary Fig. 6E-F**), highlighting its consistent performance across independent cohorts.

We next examined synaptic candidate performance across cognitive stages. ROC analyses were conducted for MCI vs Control, moderate AD vs MCI and severe vs moderate AD (**Fig. 4C**; **Supplementary Tables 3and 6**). Candidate proteins significantly discriminated each adjacent cognitive-stage comparison, although their relative performance varied across transitions. PHGDH, which mapped to the protein degradation module, achieved the highest candidate AUC for MCI vs Control and moderate AD vs MCI discrimination (AUC=0.77 and 0.67, respectively; **Fig. 5A-B**), whereas HEBP1, which mapped to CME, achieved the highest candidate AUC for severe vs moderate AD discrimination (AUC=0.85, 95% Confidence Interval [CI]=0.81-0.89, FDR=2.82×10^-27^; **Supplementary Fig. 6G**, **Supplementary Tables 3and 6**). Thus, candidates performing strongly at earlier cognitive stages were not uniformly the leading markers at later stages. These results indicate that different candidates may be informative at different cognitive stages, supporting the evaluation of marker combinations spanning distinct synaptic processes.

**Fig. 5:**
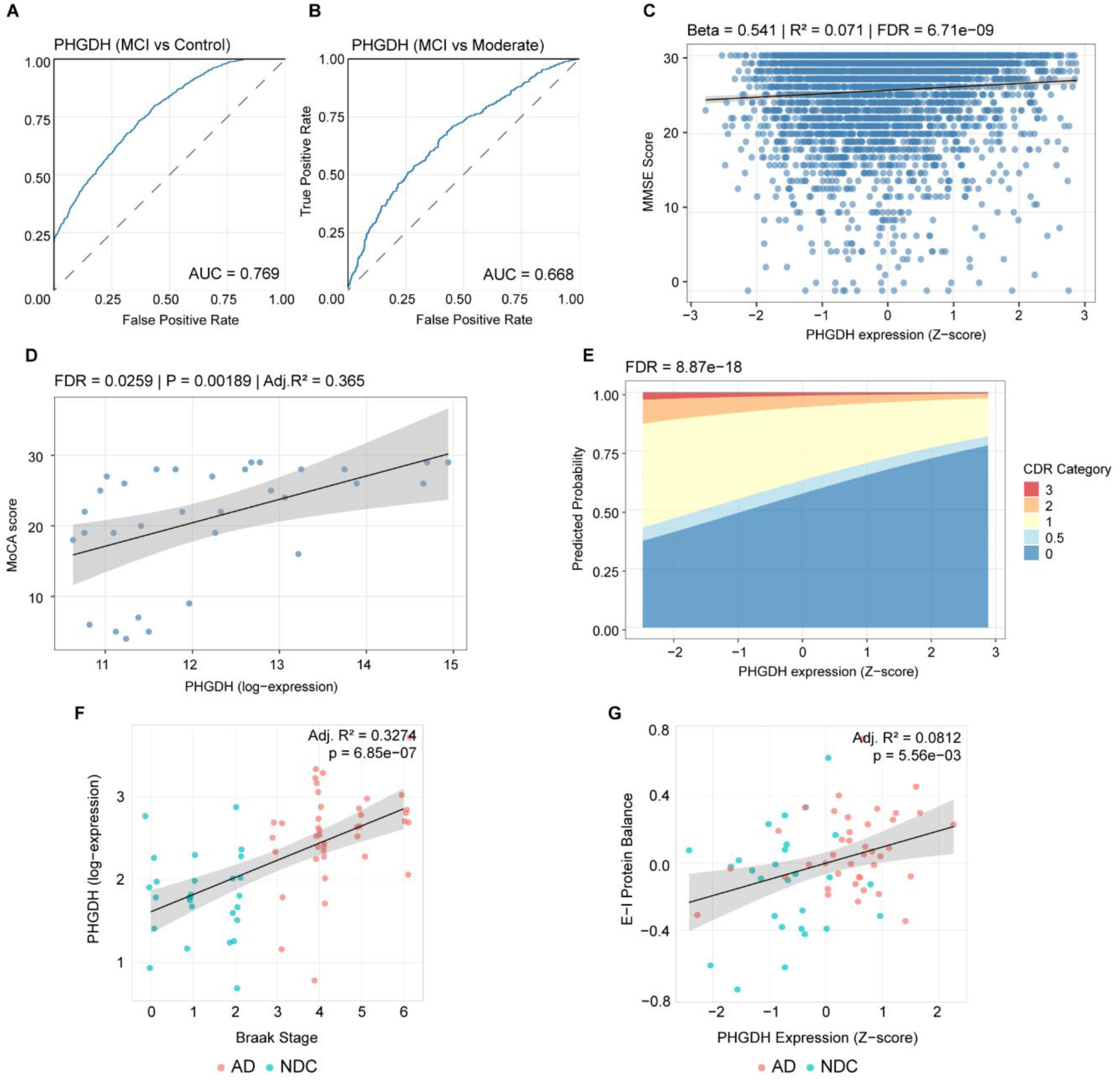
Plasma and synaptic PHGDH associate with AD severity. (**A-B**) ROC curves for plasma PHGDH discrimination of (**A**) MCI vs Control and (**B**) Moderate AD vs MCI based on cognitive examination scores in the GNPC dataset. (**C–D**) Associations of plasma PHGDH with (**C**) MMSE in GNPC and (**D**) MoCA in the Dammer et al. dataset. (**E**) Predicted probabilities of CDR categories across plasma PHGDH abundance from the partial proportional odds model in GNPC. GNPC models were adjusted for age at visit, sex, PC1 and PC2; the Dammer et al. model was adjusted for age at visit and sex. (**F**) Linear regression of PHGDH expression across Braak stages in synaptic proteomics. (**G**) Linear regression of E-I protein balance against z-scored PHGDH expression in synaptic proteomics. Models were adjusted for age, sex, PMI and batch. Lines and shaded bands show fitted linear association and 95% confidence intervals. FDR < 0.05. AD, Alzheimer’s disease; ROC, receiver operating characteristic; MCI, mild cognitive impairment; MMSE, Mini-Mental State Examination; GNPC, Global Neurodegeneration Proteomics Consortium; MoCA, Montreal Cognitive Assessment; CDR, Clinical Dementia Rating; PC, principal component; E–I, excitatory– inhibitory; PMI, post-mortem interval; FDR, false discovery rate.

To determine the ability of candidate synaptic markers to track continuous cognitive decline we performed regression analyses against cognitive examination scores (**Fig. 4D; Supplementary Tables 3and 12-14**). We conducted linear regression analyses with Mini Mental State Examination (MMSE) and Montreal Cognitive Assessment (MoCA) scores as dependent variables, using candidate protein abundance in the GNPC and Dammer et al. datasets, respectively (**Supplementary Tables 12and 13**). We additionally performed ordinal logistic regression with Clinical Dementia Rating (CDR) as the dependent variable in the GNPC dataset (**Supplementary Table 14**). Of the candidates mapped to synaptic modules, 25 proteins were significant with MMSE, 36 with MoCA and 38 with CDR. Significant associations with at least one cognitive measure were represented across all six synaptic modules, connecting the pathway-linked candidate set to continuous and ordinal measures of cognitive severity. Among these, PCSK1 and STX1A showed consistent monotonic associations with cognitive severity (**Supplementary Fig. 7**) and mapped to mitochondrial function and synaptic signalling, respectively, illustrating how cognition-associated candidates represented different synaptic signatures.

Together, the highlighted candidates linked different patterns of disease discrimination and cognitive association to distinct brain-derived synaptic signatures: Protein degradation with AD vs Control and early-stage discrimination (CNDP2, PHGDH), CME with later-stage discrimination (HEBP1), and mitochondrial function and synaptic signalling with monotonic associations with cognitive severity (PCSK1 and STX1A, respectively). These findings nominate a synaptic biomarker panel reflecting different signatures of synaptic dysfunction and providing stage-specific and cognitive trajectory information to stratify patients and inform prognostic assessment.

### PHGDH associates with neuropathological progression and cognitive severity

To illustrate how the brain-informed framework can prioritise a mechanistically interpretable candidate, we examined PHGDH, which linked several complementary analyses. Aside from being the leading candidate for early disease discrimination (**Fig. 5A-B**), PHGDH retained good discriminatory ability between severe vs moderate AD (AUC=0.84, 95% CI=0.80–0.89, FDR=5.06×10^-27^; **Supplementary Table 6**). PHGDH was also among the proteins most strongly associated with MMSE and MoCA, with lower plasma PHGDH associated with higher cognitive severity (**Fig. 5C-D**; **Supplementary Tables 12and 13**). In ordinal logistic regression, PHGDH did not track monotonic changes, but significantly captured the transition of CDR 0 (cognitively normal) to 0.5 (MCI; FDR=8.87×10^-18^, OR=1.4) consistent with its performance in the MCI vs Control analysis (**Fig. 5E**; **Supplementary Table 14**). In the synaptic proteome, PHGDH was classified as a Progression candidate and increased across Braak stages (**Fig. 5F**; **Supplementary Table 9**). Thus, brain and plasma PHGDH changed in opposite directions but were each associated with neuropathological stage or cognitive severity.

Given evidence that PHGDH promotes excitatory synaptic transmission^23^, we examined its relationship with the E–I protein balance signature in the synaptic proteomic dataset (**Fig. 5G**). PHGDH abundance increased with E–I protein balance (adjusted *P*=5.56×10^-3^), indicating that higher synaptic PHGDH expression accompanied the relative shift towards excitatory over inhibitory protein abundance across Braak stages.

PHGDH is predominantly expressed in astrocytes in the brain^24^. We therefore investigated whether the PHGDH signal captured in synaptoneurosomes could reflect the presence of perisynaptic astrocytic processes. Using immunofluorescence, we assessed the spatial association of PHGDH in ezrin^+^ perisynaptic astrocytic processes with synaptophysin^+^ synaptic puncta across AD Braak stages (**Supplementary Fig. 8A-C**). Both the PHGDH-ezrin volume fraction and the proportion of synaptophysin^+^ puncta colocalised with PHGDH-ezrin differed across stages (Kruskal–Wallis *BH-adjusted P*=0.021 and *P*=0.016, respectively) and increased with advancing pathology (Spearman *ρ*=0.775, BH-adjusted *P*=0.001 and *ρ*=0.850, BH-adjusted *P*=1.77×10^-4^, respectively). Dunn’s post-hoc analysis identified significant increases in late-stage AD compared with NDC for both measures (BH-adjusted *P*=0.011 and *P*=0.0044, respectively; **Supplementary Fig. 8D-E**). Increases were also evident at mid-stage AD but did not remain significant after correction. These results indicate stage-associated increases in PHGDH–ezrin^+^ astrocytic processes and in the proportion of synaptophysin^+^ puncta spatially associated with them.

Together, these findings position PHGDH as an illustrative candidate connecting brain, plasma and spatial analyses and showcase the mechanistic link between synaptic processes and cognitive tracking by plasma provided by the nominated candidates.

## Discussion

Cognitive decline is closely linked to synapse dysfunction in AD, yet current fluid biomarkers do not sufficiently capture the functional state of synapses^25^. Here, we took an integrated approach: First mapping the temporal trajectory of synaptic proteomic changes across AD pathology, then leveraging these mechanistic insights to identify and interpret candidate plasma markers that track distinct synaptic processes. This strategy nominated a set of plasma candidates linked to distinct synaptic pathways and disease stages, with PHGDH providing an illustrative example of how the framework mechanistically links plasma associations with stage-associated synaptic changes in the brain.

Our synaptic proteomics revealed that synaptic processes followed distinct trajectories across AD progression. Synaptic signalling and mitochondrial function declined, reaching a plateau by mid-stage AD (Braak 3-4). The shared stabilisation of these signatures is consistent with the dependence of synaptic transmission on mitochondrial ATP production and calcium buffering^26^. Synaptic vesicle recycling following neurotransmitter release depends on CME, through which clathrin-coated pits capture membrane cargo and promote vesicle budding^27,28^. Activity-dependent CME is increased at hyperactive synapses in AD and is required for Aβ production^29^, whereas impaired vesicle recycling can restrict available vesicle pools and promote the accumulation of CME-associated proteins at presynaptic terminals^30,31^. The increased CME signature at mid-stage AD may therefore capture altered endocytic demand, impaired turnover of recycling machinery or contributions from both processes. Rather than indicating continued deterioration across every pathway, the stabilisation of CME, mitochondrial function and synaptic signalling suggests that alterations in several core synaptic processes are established by mid-stage AD, before the widespread synapse loss observed in advanced disease^32^.

By contrast, proteostasis and astrocyte-associated focal adhesion increased significantly across both stage transitions. The protein degradation module, which increased more strongly at later AD stages, comprised neddylation, Cullin RING ligases and ubiquitin-proteasome pathways. Neddylation activates Cullin RING ligases, which ubiquitinate substrates for degradation, notably including tau^33^, while CUL5 expression has been associated with neuronal resilience in AD^34^. Aβ can also increase neuronal protein synthesis through MNK1–eIF4E signalling and alter the translation of synaptic proteins^35^. The parallel increase in protein synthesis and degradation signatures suggests progressive dysregulation of synaptic proteostasis with increasing AD pathology. The stronger late-stage increase in the protein degradation module and its association with pTau burden further suggest increasing demand for ubiquitin-dependent protein clearance as tau pathology accumulates.

Critically, our analysis revealed a signature of progressive increase in astrocytic engagement with synapses, demonstrated by upregulation of astrocytic focal adhesion proteins in synaptoneurosomes. The focal adhesion module is enriched in integrins (ITGA6, ITGA7, ITGB4), which are predominantly expressed in astrocytes, and their cytoskeletal anchors (filamin A, ILK)^36–38^. Their coordinated upregulation suggests enhanced binding of perisynaptic astrocytic processes (PAPs) to synaptic laminins at the extracellular matrix, effectively increasing astrocyte-synapse proximity^39^. The retraction of PAPs is crucial for learning and memory and is impaired in the APP/PS1 mouse model, consistent with our findings^40^. This evidence could point to a compensatory mechanism of an increase in reactive astrocytes associating with synapses to regulate synaptic transmission, which becomes aberrant across the pathological progression of disease.

Beyond these pathway-specific trajectories, our analysis identified a progressive shift in the relative abundance of excitatory and inhibitory synaptic proteins across AD stages. Electrophysiological measurements in human temporal cortex have shown an increased E-I current ratio in AD^22^. Protein-level findings have been less consistent: Reduced inhibitory synapse density and GABAA receptor abundance have been reported^41,42^, whereas other studies found reduced VGLUT1^+^ excitatory boutons with preservation of VGAT^43^. Proteomics in human synaptosomes also identified changes in receptor-associated proteins despite unchanged summed AMPAR/GABAAR abundance^22^. By integrating multiple markers, our signature captures coordinated relative changes without relying on individual proteins. Although both signals declined across Braak stages, the inhibitory signal declined more strongly, producing a progressive shift in E–I protein balance. Paired proteomic and electrophysiological analyses will be required to establish its relationship with altered synaptic currents.

The distinct synaptic protein signatures and their temporal dissociation reveal that AD does not uniformly disrupt all synaptic processes but alters them in a staged manner. Characterising the trajectory of each process enabled us to identify plasma candidates linked to these signatures and establish a framework for further biomarker discovery. The representation of all six synaptic modules among the 42 plasma candidates, together with variation in their associations across datasets, disease stages and measures of cognitive severity, supports their evaluation as a panel. Combining candidates linked to different synaptic processes could capture complementary aspects of synaptic dysfunction and distinguish patients characterised by alterations in different synaptic signatures. Such a panel could complement established markers of neurodegeneration for synapse-directed patient stratification and therapeutic monitoring.

PHGDH illustrates how our brain-informed framework can provide biological context for an individual plasma candidate. Although PHGDH has been previously suggested as an AD biomarker, its association as a plasma marker with synaptic changes has not been explored^44^. Synaptic proteomics showed a progressive increase of PHGDH across Braak stages. In the brain, PHGDH is primarily expressed in astrocytes and catalyses the first committed step of L-serine biosynthesis from glucose, a critical precursor for D-serine which promotes NMDAR signalling^23,24,45–47^. PHGDH inhibition reduces NMDAR potentials and hippocampal synaptic plasticity in mice, with long-term potentiation restored by exogenous L-serine^23^. Our spatial analyses showed increases in PHGDH-ezrin^+^ PAPs and in the proportion of synaptophysin^+^ puncta associated with them across AD stages, supporting a perisynaptic astrocytic contribution to the PHGDH captured in synaptoneurosomes. PHGDH abundance was also associated with the E–I protein balance signature, consistent with its established role in promoting excitatory synaptic transmission^23,48^. PHGDH mapped to the protein degradation module, which consistently increased in association to pTau and Aβ accumulation. Recent experimental evidence linked PHGDH to altered protein clearance and increased Aβ production via a mechanism independent of its enzymatic activity, whereby PHGDH promoted astrocytic transcription of IKKα and HMGB1^49^. To test its translational validity, we showed that plasma PHGDH discriminated across cognitive stages and declined with increased cognitive severity assessed by MMSE, MoCA and CDR. Lower plasma PHGDH has similarly been associated with greater cerebral β-amyloidosis in AD^50^, while PHGDH is also downregulated in CSF in AD^13^. This discordance between brain and fluid profiles could reflect compartment-specific responses to a shared disease-associated metabolic process or regulated transport of glycolytic proteins across the blood–brain barrier^50^. Together, PHGDH illustrates how our framework connects plasma associations with stage-associated molecular and spatial synaptic changes, providing biological context for individual members of the broader pathway-informed candidate set. Further functional and longitudinal studies will be required to establish causality and evaluate their clinical utility.

The use of human tissue in our study enabled the neuropathological confirmation of AD alongside a clinical diagnosis, thus providing a relevant and reliable disease paradigm for a comprehensive characterisation of synaptic alterations in disease. To ensure the robustness of our findings, we performed orthogonal validation using independent public datasets of plasma proteomics via different platforms, as well as alternative definitions of disease progression based on cognitive decline and not solely on pathology. Therefore, we report results that can be translatable in the clinic alongside a biological interpretation of synaptic dysfunction. However, we recognise inherent limitations of our research. First, *post-mortem* human brain tissue reflects the disease context in humans but may be influenced by variability in tissue quality. This was mitigated by including PMI as a covariate in our models. Importantly, the findings reported here are correlational and further functional studies are required to establish causal relationships in the mechanisms proposed in our study. Furthermore, as exhibited both by our data and the literature, the Syn-PER method of synaptoneurosome isolation is prone to contamination by other cell types^51^. However, this preparation method allows for a more comprehensive study of synapses by preserving the post-synaptic compartment and maintaining interactions with other cells that associate with synapses in the brain. Despite these limitations, our results open new avenues for discovery of biomarkers tracking synaptopathy in AD progression, as well as novel targets for synaptotherapeutic drugs.

In conclusion, our study identified proteomic markers of distinct processes of synaptic dysfunction which can track cognitive impairment when detected in plasma. Despite fluid biomarkers of synaptic dysfunction having previously been proposed, there has been little understanding regarding the biological mechanisms associated with their alterations in disease. We conduct an unbiased comprehensive characterisation of the synaptic proteome across AD and later perform targeted investigation of candidate synaptic biomarkers in biofluids. We provide a panel of synaptic plasma candidates that can differentially discriminate across different stages of the disease and track cognitive decline. Therefore, our results could have useful implications not only in progression tracking, but also patient stratification and disease staging. Finally, the introduction of biomarkers indicating the state of synaptic health could not only facilitate drug development by providing useful, easily accessible methods of measuring the impact of a drug on synaptic dysfunction but may also point to novel personalised therapeutic targets to restore synaptic function in adjunct to disease modifying therapies.

## Data availability

All data and code associated with this manuscript will be available after peer review.

## Funding

This work was supported by funding from Alzheimer’s Society (grant number 599 (AS-DRL-22-008)) which also supports JSJ and TG. Funding was also received for this work from UK Dementia Research Institute, which receives funding from UK DRI Ltd., funded by the UK Medical Research Council, Alzheimer’s Society, and Alzheimer’s Research UK which also supported JG. PMM acknowledges generous personal support from the Edmond J Safra Foundation and Lily Safra and an NIHR Senior Investigator Award. The study is an output from the UK Dementia Institute Multi-omics Atlas Project for Alzheimer’s Disease (MAP- AD; map-ad.org). Infrastructure, including particularly the LMS/NIHR Imperial Biomedical Research Centre Flow Cytometry Facility and the Imperial BRC Genomics Facility, was supported by the National Institute for Health Research (NIHR) Biomedical Research Centre (BRC) (grant number MC_PC_MR/X013537/1)

Work in the SG laboratory was funded by the UK Dementia Research Institute which receives its funding from UK DRI Ltd (award XCPD2019–01) funded by the UK Medical Research Council (MR/X021564/1), Chan Zuckerberg Initiative (DAF2025-351071) and Alzheimer’s Society and Alzheimer’s Research UK (JG).

## Acknowledgements

We thank the donors and their families for the use of human brain tissue in this study and the UK brain bank staff for making it available. Tissue samples were provided by the London Neurodegenerative Diseases Brain Bank at King’s College London. The brain bank receives funding from the UK Medical Research Council and as part of the Brains for Dementia Research programme, jointly funded by Alzheimer’s Research UK and the Alzheimer’s Society. We would like to thank the South West Dementia Brain Bank (SWDBB) for providing brain tissue for this study. The SWDBB is part of the Brains for Dementia Research programme, jointly funded by Alzheimer’s Research UK and Alzheimer’s Society and is supported by BRACE (Bristol Research into Alzheimer’s and Care of the Elderly) and the Medical Research Council. We also thank the Edinburgh Brain Bank for the provision of tissue and expert advice from neuropathologist, Professor Colin Smith. The provision of case data used in this study was provided with support from the BDR programme, jointly funded by Alzheimer’s Society UK and Alzheimer’s Society. We are grateful to Diana P. Benitez for her support in the human tissue management.

This work was supported by funding from Alzheimer’s Society (grant number 599 (AS-DRL-22-008)) which also supports JSJ and TG. Funding was also received for this work from UK Dementia Research Institute, which receives funding from UK DRI Ltd., funded by the UK Medical Research Council, Alzheimer’s Society, and Alzheimer’s Research UK which also supported JG. PMM acknowledges generous personal support from the Edmond J Safra Foundation and Lily Safra and an NIHR Senior Investigator Award. The study is an output from the UK Dementia Institute Multi-omics Atlas Project for Alzheimer’s Disease (MAP-AD; map-ad.org). Infrastructure, including particularly the LMS/NIHR Imperial Biomedical Research Centre Flow Cytometry Facility and the Imperial BRC Genomics Facility, was supported by the National Institute for Health Research (NIHR) Biomedical Research Centre (BRC) (grant number MC_PC_MR/X013537/1)

## Author contributions

TG conceived and designed the study, performed wet-lab experiments, analysed data and drafted the manuscript. JG performed sample preparation of the late-stage cohort. FG contributed to data analysis and literature review. EC and MP performed immunohistochemistry experiments. DL performed APOE genotyping. HW performed LC– MS analyses. PMM, SJB and SG contributed to study guidance, supervision and funding acquisition. JSJ conceived and designed the study, supervised the project, acquired funding and revised the manuscript. All authors reviewed the manuscript, provided critical feedback and approved the final version.

## Notes

https://www.nature.com/articles/s41591-025-03834-0

https://link.springer.com/article/10.1186/s13195-022-01113-5

### Competing Interest Statement

The authors have declared no competing interest.

## References

1. Griffiths J, Schneegans E, Whitwell H, et al. A synaptic-astrocytic proteomic signature associated with synaptopathy in Alzheimer’s Disease. bioRxiv. Preprint posted online January 23, 2025:2025.01.23.634408. doi:10.1101/2025.01.23.634408

2. Gatto L, Gibb S, Rainer J. MSnbase, Efficient and Elegant R-Based Processing and Visualization of Raw Mass Spectrometry Data. J Proteome Res. 2021;20(1):1063–1069. doi:10.1021/acs.jproteome.0c00313

3. Ritchie ME, Phipson B, Wu D, et al. limma powers differential expression analyses for RNA-sequencing and microarray studies. Nucleic Acids Res. 2015;43(7):e47. doi:10.1093/nar/gkv007

4. Benjamini Y, Hochberg Y. Controlling the False Discovery Rate: A Practical and Powerful Approach to Multiple Testing. Journal of the Royal Statistical Society: Series B (Methodological). 1995;57(1):289–300. doi:10.1111/j.2517-6161.1995.tb02031.x

5. Langfelder P, Horvath S. WGCNA: an R package for weighted correlation network analysis. BMC Bioinformatics. 2008;9(1):559. doi:10.1186/1471-2105-9-559

6. Kuleshov MV, Jones MR, Rouillard AD, et al. Enrichr: a comprehensive gene set enrichment analysis web server 2016 update. Nucleic Acids Res. 2016;44(Web Server issue):W90–W97. doi:10.1093/nar/gkw377

7. Skene NG, Grant SGN. Identification of Vulnerable Cell Types in Major Brain Disorders Using Single Cell Transcriptomes and Expression Weighted Cell Type Enrichment. Front Neurosci. 2016;10. doi:10.3389/fnins.2016.00016

8. Hodge RD, Bakken TE, Miller JA, et al. Conserved cell types with divergent features in human versus mouse cortex. Nature. 2019;573(7772):61–68. doi:10.1038/s41586-019-1506-7

9. Hänzelmann S, Castelo R, Guinney J. GSVA: gene set variation analysis for microarray and RNA-Seq data. BMC Bioinformatics. 2013;14(1):7. doi:10.1186/1471-2105-14-7

10. Lauterborn JC, Scaduto P, Cox CD, et al. Increased excitatory to inhibitory synaptic ratio in parietal cortex samples from individuals with Alzheimer’s disease. Nat Commun. 2021;12(1):2603. doi:10.1038/s41467-021-22742-8

11. Imam F, Saloner R, Vogel JW, et al. The Global Neurodegeneration Proteomics Consortium: biomarker and drug target discovery for common neurodegenerative diseases and aging. Nat Med. 2025;31(8):2556–2566. doi:10.1038/s41591-025-03834-0

12. Ali M, Erabadda B, Chen Y, et al. Shared and disease-specific pathways in frontotemporal dementia and Alzheimer’s and Parkinson’s diseases. Nat Med. 2025;31(8):2567–2577. doi:10.1038/s41591-025-03833-1

13. Dammer EB, Ping L, Duong DM, et al. Multi-platform proteomic analysis of Alzheimer’s disease cerebrospinal fluid and plasma reveals network biomarkers associated with proteostasis and the matrisome. Alzheimer’s Research & Therapy. 2022;14(1):174. doi:10.1186/s13195-022-01113-5

14. Perneczky R, Wagenpfeil S, Komossa K, Grimmer T, Diehl J, Kurz A. Mapping Scores Onto Stages: Mini-Mental State Examination and Clinical Dementia Rating. The American Journal of Geriatric Psychiatry. 2006;14(2):139–144. doi:10.1097/01.JGP.0000192478.82189.a8

15. Nasreddine ZS, Phillips NA, Bédirian V, et al. The Montreal Cognitive Assessment, MoCA: A Brief Screening Tool For Mild Cognitive Impairment. Journal of the American Geriatrics Society. 2005;53(4):695–699. doi:10.1111/j.1532-5415.2005.53221.x

16. Christensen RHB. Ordinal—Regression Models for Ordinal Data. 2026. https://CRAN.R-project.org/package=ordinal

17. Nguyen P, Zeng E. A Protocol for Weighted Gene Co-expression Network Analysis With Module Preservation and Functional Enrichment Analysis for Tumor and Normal Transcriptomic Data. Bio Protoc. 2025;15(18):e5447. doi:10.21769/BioProtoc.5447

18. Yang C, Colosi P, Hugelier S, Zabezhinsky D, Lakadamyali M, Svitkina T. Actin polymerization promotes invagination of flat clathrin-coated lattices in mammalian cells by pushing at lattice edges. Nature Communications. 2022;13(1):6127. doi:10.1038/s41467-022-33852-2

19. Escartin C, Galea E, Lakatos A, et al. Reactive astrocyte nomenclature, definitions, and future directions. Nat Neurosci. 2021;24(3):312–325. doi:10.1038/s41593-020-00783-4

20. Al-Dalahmah O, Sosunov AA, Shaik A, et al. Single-nucleus RNA-seq identifies Huntington disease astrocyte states. Acta Neuropathol Commun. 2020;8:19. doi:10.1186/s40478-020-0880-6

21. Mathys H, Davila-Velderrain J, Peng Z, et al. Single-cell transcriptomic analysis of Alzheimer’s disease. Nature. 2019;570(7761):332–337. doi:10.1038/s41586-019-1195-2

22. Scaduto P, Lauterborn JC, Cox CD, et al. Functional excitatory to inhibitory synaptic imbalance and loss of cognitive performance in people with Alzheimer’s disease neuropathologic change. Acta Neuropathologica. 2023;145(3):303–324. doi:10.1007/s00401-022-02526-0

23. Neame S, Safory H, Radzishevsky I, et al. The NMDA receptor activation by d-serine and glycine is controlled by an astrocytic Phgdh-dependent serine shuttle. Proc Natl Acad Sci U S A. 2019;116(41):20736–20742. doi:10.1073/pnas.1909458116

24. Hawrylycz M, Kaplan ES, Travaglini KJ, et al. SEA-AD: A multimodal cellular atlas and resource for Alzheimer’s disease. Nat Aging. 2024;4(10):1331–1334. doi:10.1038/s43587-024-00719-8

25. Bereczki E, Branca RM, Francis PT, et al. Synaptic markers of cognitive decline in neurodegenerative diseases: a proteomic approach. Brain. 2018;141(2):582–595. doi:10.1093/brain/awx352

26. Cai Q, Tammineni P. Mitochondrial Aspects of Synaptic Dysfunction in Alzheimer’s Disease. J Alzheimers Dis. 2017;57(4):1087–1103. doi:10.3233/JAD-160726

27. Royle SJ, Lagnado L. Clathrin-mediated endocytosis at the synaptic terminal: bridging the gap between physiology and molecules. Traffic. 2010;11(12):1489–1497. doi:10.1111/j.1600-0854.2010.01104.x

28. Milosevic I. Revisiting the Role of Clathrin-Mediated Endoytosis in Synaptic Vesicle Recycling. Front Cell Neurosci. 2018;12. doi:10.3389/fncel.2018.00027

29. Stargardt A, Swaab DF, Bossers K. The storm before the quiet: neuronal hyperactivity and Aβ in the presymptomatic stages of Alzheimer’s disease. Neurobiology of Aging. 2015;36(1):1–11. doi:10.1016/j.neurobiolaging.2014.08.014

30. Jaye S, Sandau US, Saugstad JA. Clathrin mediated endocytosis in Alzheimer’s disease: cell type specific involvement in amyloid beta pathology. Front Aging Neurosci. 2024;16:1378576. doi:10.3389/fnagi.2024.1378576

31. Alabi AA, Tsien RW. Synaptic Vesicle Pools and Dynamics. Cold Spring Harb Perspect Biol. 2012;4(8):a013680. doi:10.1101/cshperspect.a013680

32. Taddei RN, E. Duff K. Synapse vulnerability and resilience underlying Alzheimer’s disease. eBioMedicine. 2025;112:105557. doi:10.1016/j.ebiom.2025.105557

33. Zhang S, Yu Q, Li Z, Zhao Y, Sun Y. Protein neddylation and its role in health and diseases. Signal Transduction and Targeted Therapy. 2024;9(1):85. doi:10.1038/s41392-024-01800-9

34. Samelson AJ, Ariqat N, McKetney J, et al. CRISPR screens in iPSC-derived neurons reveal principles of tau proteostasis. Cell. 2026;189(5):1517–1534.e19. doi:10.1016/j.cell.2025.12.038

35. Ghosh A, Mizuno K, Tiwari SS, et al. Alzheimer’s disease-related dysregulation of mRNA translation causes key pathological features with ageing. Transl Psychiatry. 2020;10(1):192. doi:10.1038/s41398-020-00882-7

36. Gabitto MI, Travaglini KJ, Rachleff VM, et al. Integrated multimodal cell atlas of Alzheimer’s disease. Nat Neurosci. 2024;27(12):2366–2383. doi:10.1038/s41593-024-01774-5

37. Golushko NI, Shevlyakov AD, Martynov DD, et al. The Emerging Role of Filamin A and FLNA Gene in Central Nervous System Functions: Insights into Neurodevelopment and Disease. J Integr Neurosci. 2025;24(6). doi:10.31083/JIN26489

38. Qin J, Wu C. ILK: a pseudokinase in the center stage of cell-matrix adhesion and signaling. Current Opinion in Cell Biology. 2012;24(5):607–613. doi:10.1016/j.ceb.2012.06.003

39. Pérez LA, Rashid A, Combs JD, et al. An Outside-In Switch in Integrin Signaling Caused by Chemical and Mechanical Signals in Reactive Astrocytes. Front Cell Dev Biol. 2021;9:712627. doi:10.3389/fcell.2021.712627

40. Kater MSJ, Badia-Soteras A, van Weering JRT, Smit AB, Verheijen MHG. Electron microscopy analysis of astrocyte-synapse interactions shows altered dynamics in an Alzheimer’s disease mouse model. Front Cell Neurosci. 2023;17:1085690. doi:10.3389/fncel.2023.1085690

41. Kurucu H, Colom-Cadena M, Davies C, et al. Inhibitory synapse loss and accumulation of amyloid beta in inhibitory presynaptic terminals in Alzheimer’s disease. European Journal of Neurology. 2022;29(5):1311–1323. doi:10.1111/ene.15043

42. Limon A, Reyes-Ruiz JM, Miledi R. Loss of functional GABAA receptors in the Alzheimer diseased brain. Proc Natl Acad Sci U S A. 2012;109(25):10071–10076. doi:10.1073/pnas.1204606109

43. Mitew S, Kirkcaldie MTK, Dickson TC, Vickers JC. Altered synapses and gliotransmission in Alzheimer’s disease and AD model mice. Neurobiology of Aging. 2013;34(10):2341–2351. doi:10.1016/j.neurobiolaging.2013.04.010

44. Yan Z, Zhou Z, Wu Q, Chen ZB, Koo EH, Zhong S. Presymptomatic Increase of an Extracellular RNA in Blood Plasma Associates with the Development of Alzheimer’s Disease. Current Biology. 2020;30(10):1771–1782.e3. doi:10.1016/j.cub.2020.02.084

45. de Miranda J, Panizzutti R, Foltyn VN, Wolosker H. Cofactors of serine racemase that physiologically stimulate the synthesis of the N-methyl-d-aspartate (NMDA) receptor coagonist d-serine. Proceedings of the National Academy of Sciences. 2002;99(22):14542–14547. doi:10.1073/pnas.222421299

46. Yamasaki M, Yamada K, Furuya S, Mitoma J, Hirabayashi Y, Watanabe M. 3-Phosphoglycerate Dehydrogenase, a Key Enzyme forl-Serine Biosynthesis, Is Preferentially Expressed in the Radial Glia/Astrocyte Lineage and Olfactory Ensheathing Glia in the Mouse Brain. J Neurosci. 2001;21(19):7691–7704. doi:10.1523/JNEUROSCI.21-19-07691.2001

47. Basu AC, Tsai GE, Ma CL, et al. Targeted disruption of serine racemase affects glutamatergic neurotransmission and behavior. Mol Psychiatry. 2009;14(7):719–727. doi:10.1038/mp.2008.130

48. Orzylowski M, Fujiwara E, Mousseau DD, Baker GB. An Overview of the Involvement of D-Serine in Cognitive Impairment in Normal Aging and Dementia. Front Psychiatry. 2021;12. doi:10.3389/fpsyt.2021.754032

49. Chen J, Hadi F, Wen X, et al. Transcriptional regulation by PHGDH drives amyloid pathology in Alzheimer’s disease. Cell. 2025;188(13):3513–3529.e26. doi:10.1016/j.cell.2025.03.045

50. Afshar S, Dammer EB, Bian S, et al. Plasma proteomic associations with Alzheimer’s disease endophenotypes. Nat Aging. 2025;5(10):2104–2124. doi:10.1038/s43587-025-00965-4

51. Gulyássy P, Puska G, Györffy BA, et al. Proteomic comparison of different synaptosome preparation procedures. Amino Acids. 2020;52(11):1529–1543. doi:10.1007/s00726-020-02912-6

