## Supplementary Figures for "Stage-specific proteomic signatures of synaptopathy in Alzheimer’s disease"

Sir Michael Uren Hub, 86 Wood Ln, London W12 0BZ, UK

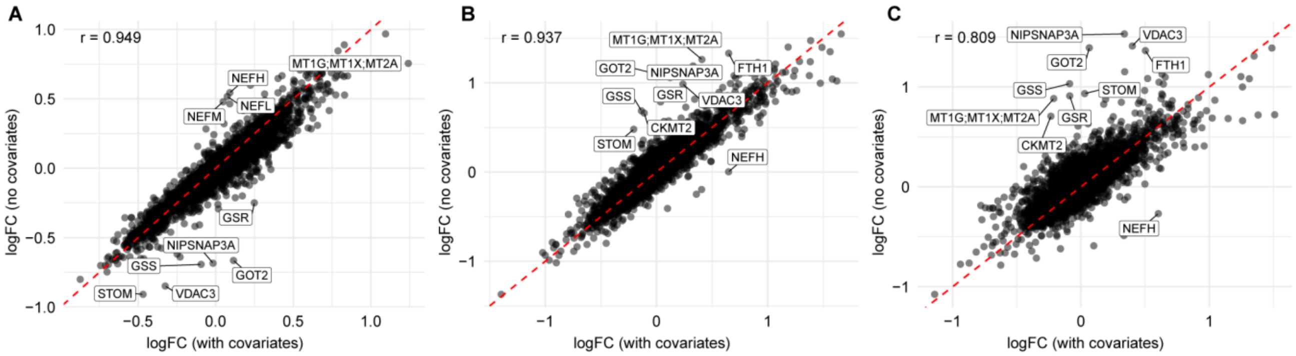


**Supplementary Fig. 1 : Covariate sensitivity analysis of differential protein expression.** (**A-C**) Correlations between log2 fold-change estimates from models fitted with and without covariate adjustment for (**A**) mid-stage AD vs NDC, (**B**) late-stage AD vs NDC and (**C**) Late vs Moderate AD. Covariate-adjusted models included age, sex, *post-mortem* interval and batch. The ten proteins showing the greatest deviation from the identity line are labelled. AD, Alzheimer's disease; NDC, non-diseased control.

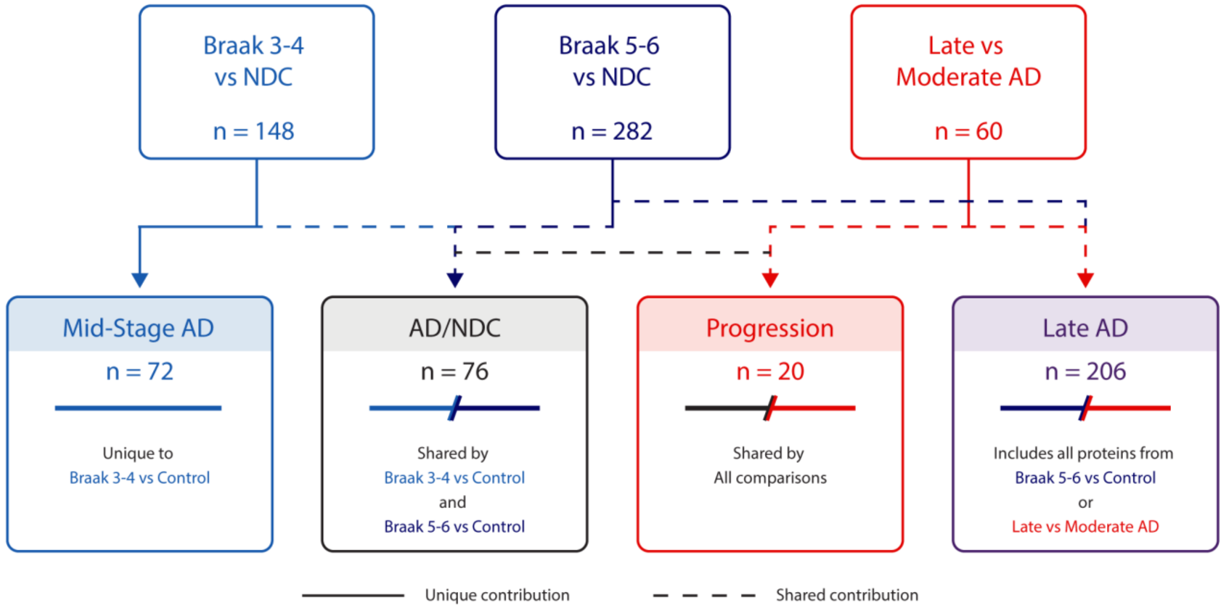


**Supplementary Fig. 2: Classification of stage-associated DEPs for plasma candidate discovery.** DEPs were classified as Mid-stage AD when unique to Braak 3–4 vs NDC, Late AD when detected in Braak 5–6 vs NDC or Late vs Moderate AD but not mid-stage AD, and AD/NDC when shared between the two comparisons with NDC. The 20 proteins significant across all three comparisons were additionally classified as Progression proteins. Final filtered counts were 72 Mid-stage AD, 206 Late AD, 76 AD/NDC and 20 Progression proteins. NDC, non-diseased control; AD, Alzheimer's disease; DEPs, differentially expressed proteins.

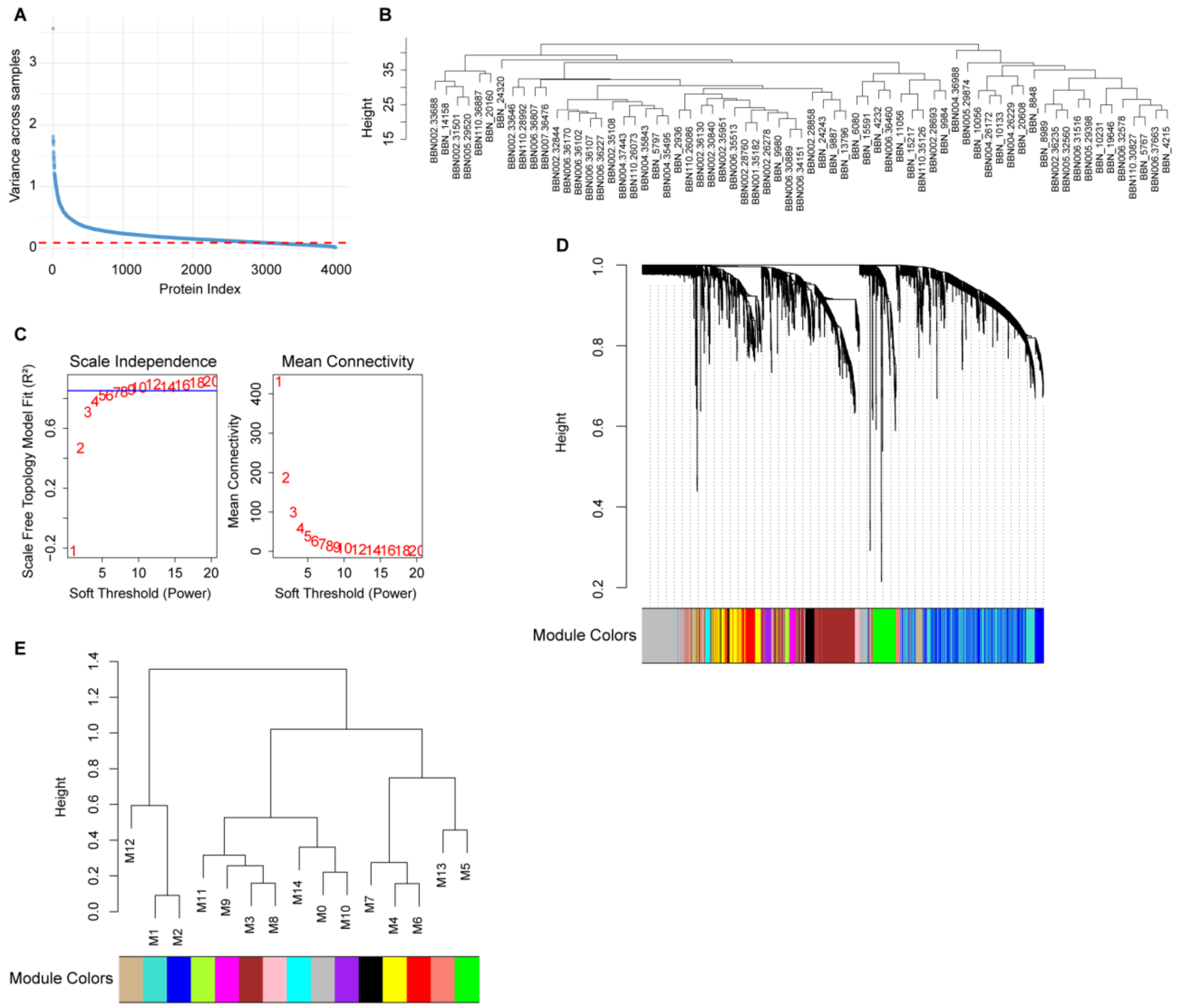


**Supplementary Fig. 3: WGCNA modules plots and quality control.** (**A**) Protein variance across samples. Red line indicates the 75th percentile of protein variance, where proteins were filtered for WGCNA. (**B**) Samples clustering tree to detect outliers. (**C**) Scale independence and mean connectivity plots were used to select the optimal soft thresholding power. Blue line indicates the 0.85 cut-off used. (**D**) Hierarchical gene clustering tree was cut at default 0.998 height to classify genes into modules. Module colours correspond to different modules mapped in (**E**). (**E**) Hierarchical clustering of modules reveals which modules are more closely related. Each module is mapped to a colour assigned by the WGCNA package and consists of genes clustering together as shown in panel (**D**). WGCNA, Weighted gene co-expression network analysis.

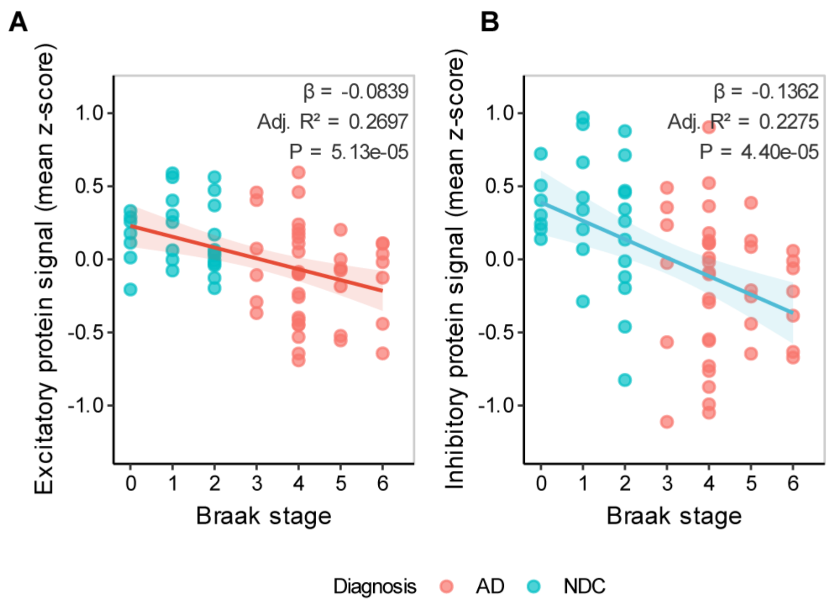


**Supplementary Fig. 4: E-I protein balance components are downregulated across AD progression.** (**A**) Associations of the mean z-scored abundance of proteins classified by Lauterborn et al. as (**A**) excitatory or (**B**) inhibitory with Braak stage across 67 synaptoneurosome samples. Models were adjusted for age, sex, *post-mortem* interval and batch. Lines and shaded bands show the fitted linear associations and 95% confidence intervals. E-I, excitatory-inhibitory.

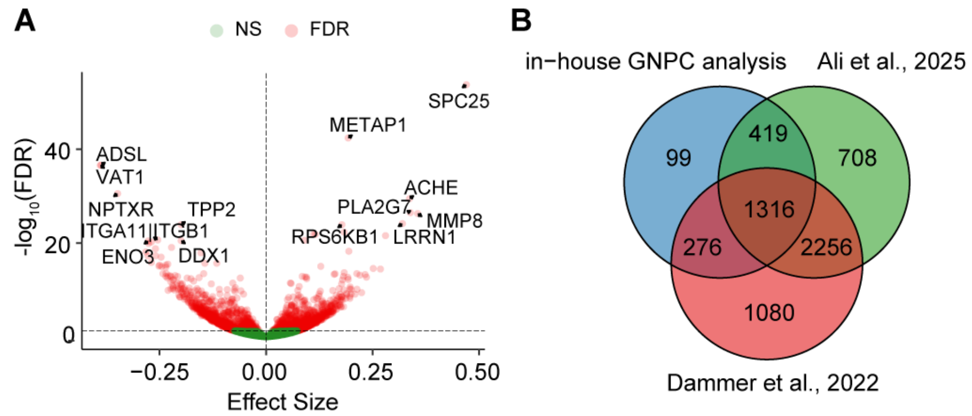


**Supplementary Fig. 5: Plasma biomarker datasets differential expression.** (**A**) Volcano plot of differential expression analysis of plasma SomaScan 7K platform proteomics in AD compared to Control cases using the GNPC dataset. Top 7 upregulated and downregulated proteins are labelled. FDR < 0.05. Linear model adjusted for age at visit, sex, PC1 and PC2. (**B**) Venn diagram of the overlap of DEPs between our in-house GNPC differential expression analysis, the analysis of the same dataset by Ali et al., 2025 and the independent Dammer et al., 2022 dataset. AD, Alzheimer's disease; GNPC, Global Neurodegeneration Proteomics Consortium; FDR, false discovery rate; PC, principal component; DEP, differentially expressed protein.

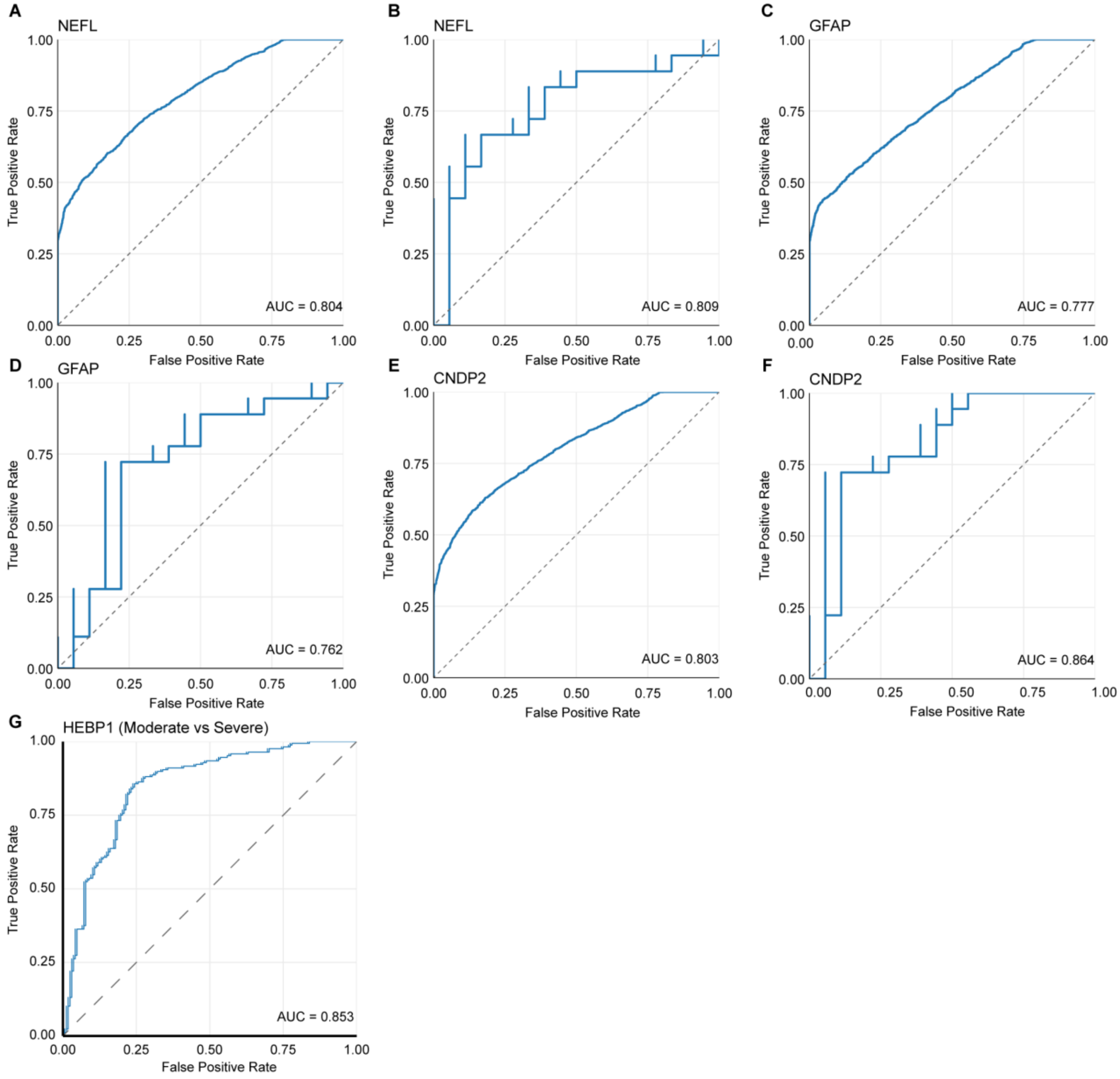


**Supplementary Fig. 6: ROC analyses of selected plasma candidates and neurodegeneration-associated comparators.** (**A-B**) NEFL discrimination of AD vs Control in (**A**) GNPC and (**B**) the Dammer et al. dataset. (**C-D**) GFAP discrimination of AD vs Control in (**C**) GNPC and (**D**) the Dammer et al. dataset. (**E-F**) CNDP2 discrimination of AD vs Control in (**E**) GNPC and (**F**) the Dammer et al. dataset. (**G**) HEBP1 discrimination of cognitively severe vs moderate AD cases in GNPC. ROC, receiver operating characteristic; AD, Alzheimer's disease; GNPC, Global Neurodegeneration Proteomics Consortium;  AUC= Area Under Curve.

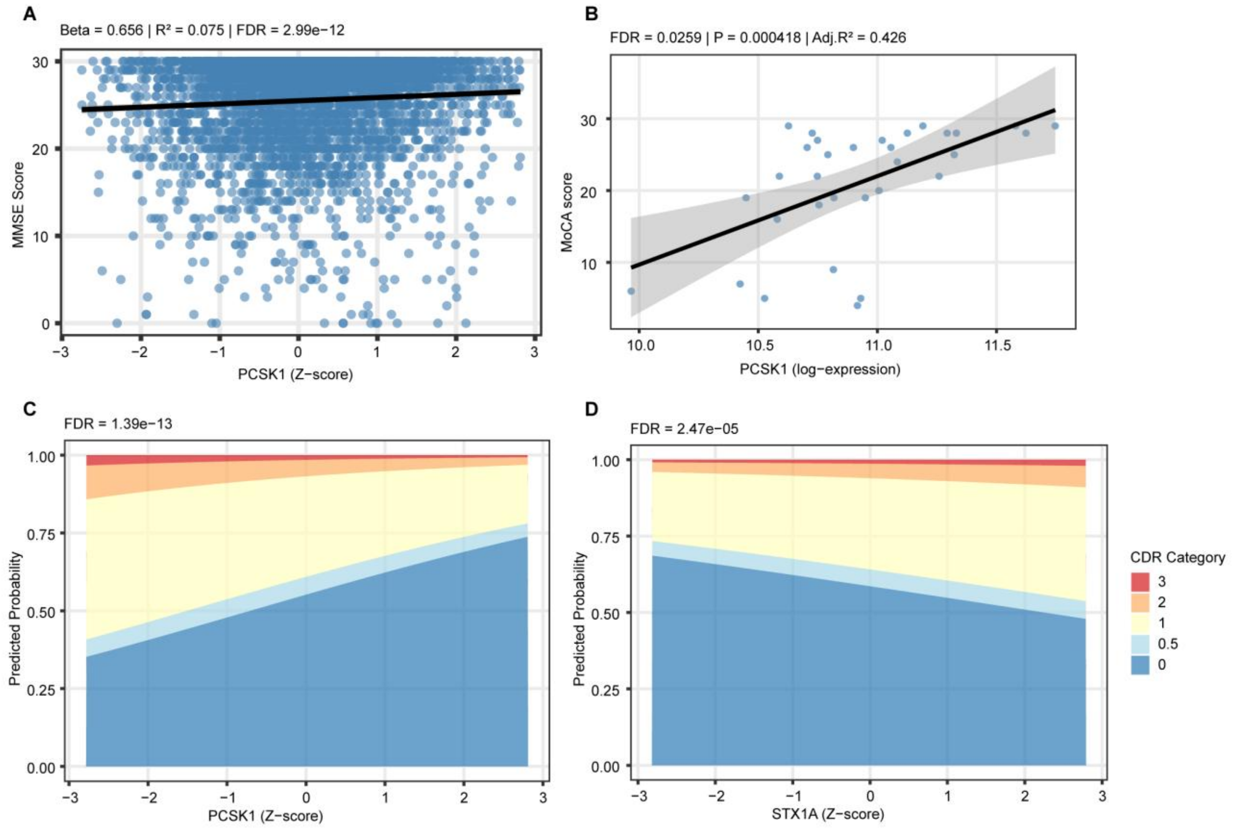


**Supplementary Fig. 7: Associations of selected plasma candidates with cognitive severity.** (**A**) Linear regression plot of plasma PCSK1 protein expression against MMSE scores in AD and Control cases using the GNPC dataset. (**B**) Linear regression plots of plasma PCSK1 protein expression against MoCA scores in AD and Control cases using the Dammer et al. dataset. Lines and shaded bands show the fitted linear associations and 95% confidence intervals. (**C-D**) Predicted probability plots of CDR severity based on plasma protein expression of (**C**) PCSK1 and (**D**) STX1A, calculated using odds ratios derived from ordinal logistic regression using the GNPC dataset. GNPC models were adjusted for age at visit, sex, PC1 and PC2; the Dammer et al. model was adjusted for age at visit and sex. MMSE, Mini-Mental State Examination; AD, Alzheimer's disease; GNPC, Global Neurodegeneration Proteomics Consortium; MoCA, Montreal Cognitive Assessment; CDR, Clinical Dementia Rating; PC, principal component.

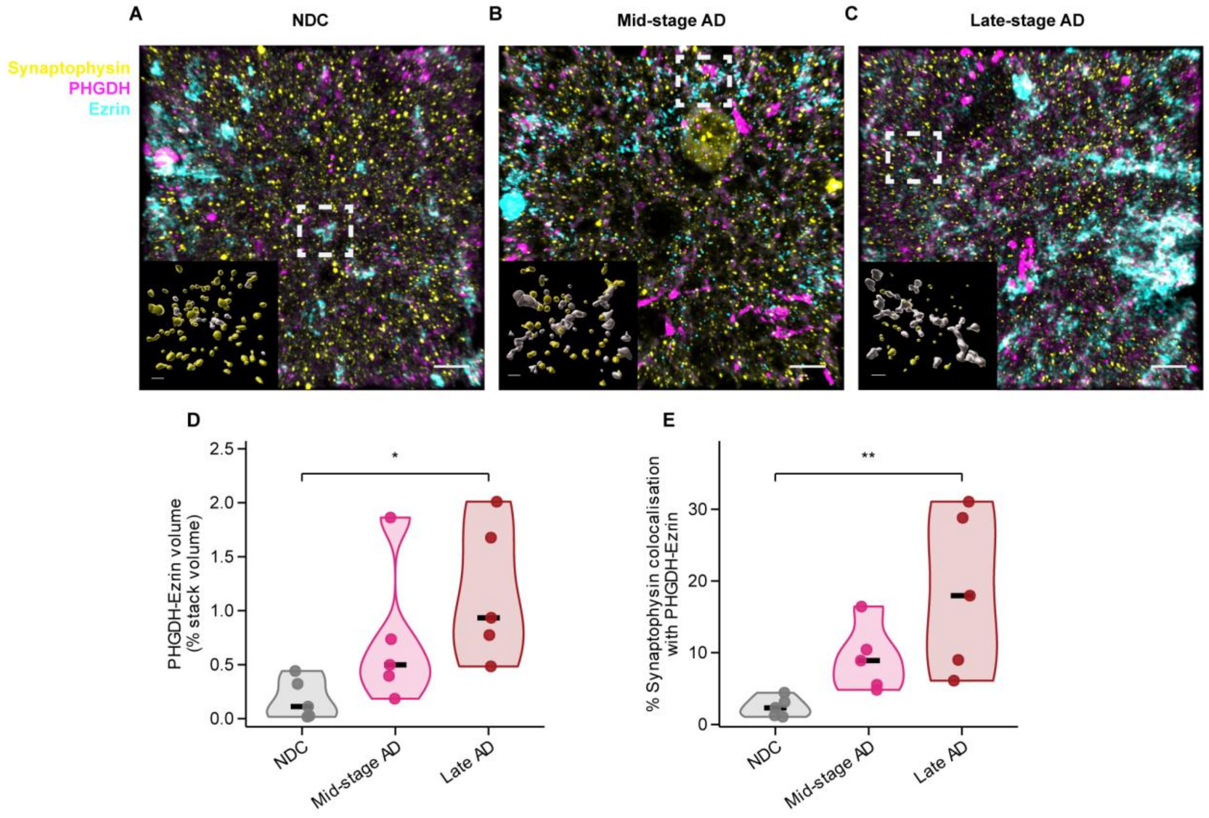


**Supplementary Fig. 8: PHGDH–ezrin⁺ astrocytic processes increasingly associate with synaptophysin puncta across AD stages.** (**A-C**) Representative Imaris 3D renderings of z-stacks from MTG FFPE sections stained for synaptophysin (yellow), PHGDH (magenta) and ezrin (cyan) from (**A**) NDC, (**B**), mid-stage AD and (**C**), late-stage AD cases. Main-image scale bars, 5 µm. Insets show surface renderings of synaptophysin$^{+}$ puncta (yellow) and PHGDH–ezrin$^{+}$ surfaces (white); inset scale bars, 1 µm. (**D**) PHGDH–ezrin$^{+}$ surface volume as a percentage of total image-stack volume. (**E**) Percentage of synaptophysin$^{+}$ puncta colocalised with PHGDH–ezrin$^{+}$ surfaces, defined by shortest distance to the surface of ≤0 µm. Each point represents the mean of three regions of interest from one case ($n = 5$ cases per group). Group differences were assessed using Kruskal–Wallis tests followed by Dunn's post-hoc tests. *P* values were adjusted using the BH method. *$P < 0.05$; **$P < 0.01$. AD, Alzheimer's disease; MTG, Middle Temporal Gyrus; FFPE, Formalin-Fixed Paraffin-Embedded.
